# Ecological theory of mutualism: Models generalizing across different mechanisms

**DOI:** 10.1101/2020.10.25.343087

**Authors:** Kayla R. S. Hale, Daniel P. Maes, Fernanda S. Valdovinos

## Abstract

Mutualisms are ubiquitous in nature, provide important ecosystem services, and involve many species of interest for conservation. Theoretical progress on the population dynamics of mutualistic interactions, however, has comparatively lagged behind that of trophic and competitive interactions. Consequently, ecologists still lack a generalized framework to investigate the population dynamics of mutualisms. Here, we propose extensible models for two-species mutualisms focusing on nutritional, protection, and transportation mechanisms and evaluate the population-level consequences of those mechanisms. We introduce a novel theoretical framework that highlights characteristic dynamics when the effects of mutualism are directly dependent or independent of recipient density and when they saturate due to inter- or intra-specific density-dependence. We end by integrating our work into the broader historical context of population-dynamic models of mutualism and conclude that a general ecological theory of mutualism exists.

## Iantroduction

Mutually beneficial interactions are ubiquitous in nature. Nearly all species on Earth participate in at least one of four main types of mutualism: seed dispersal, pollination, protection, and resource exchange including with symbionts (Janzen 1985, Bronstein 2015a,b). These interactions also support an immense amount of ecosystem function. For example, up to ~3/4 of phosphorus and nitrogen acquired by plants is provided by mycorrhizal fungi and nitrogen-fixing bacteria (van der Heijden *et al.* 2008) and ~1/3 of crop production is dependent on animal pollination (Klein *et al*. 2007). Yet, the importance of mutualism for ecological communities has only recently been recognized (Bronstein 2015b) and, accordingly, theory has lagged behind. Such theory is critical as anthropogenic perturbations like climate change, nutrient runoff, pesticide use, and invasive species increasingly threaten many mutualisms and the ecosystem services they provide (Stachowicz 2001, Tylianakis *et al.* 2008). Here, we develop simple but extensible theory (i.e., can be specified to particular systems or generalized to networks of species interactions) on the population dynamics of two-species mutualisms, which integrates mutualisms into the broader framework of community ecology.

Theoretical study of mutualism has lagged behind the other two “pillars” of community ecology: competition and predator-prey interactions (Callaway 2007). This has been attributed to many reasons, three of which we highlight here. First, the terms used to identify interactions as “mutualism” have changed over time. Previous theory treated mutualism as a subset of facilitation, in which one species alters the environment to benefit a neighboring species (Callaway 2007), or symbiosis, in which species coexist in “prolonged physical intimacy” (Bronstein 2015b), or used those terms interchangeably. We limit our scope to mutualism defined as reciprocally beneficial interactions between species without reference to the partners’ intimacy or environmental effects, though our work could be adapted to these other cases (see, e.g., Thompson *et al.* 2006).

Second, incredible diversity among mutualisms led researchers to focus on natural history, resulting in a heterogeneous set of case studies with little to no theory to unify or partition them (Addicott 1981, Bronstein 2015b). Some conceptual frameworks have attempted to organize this rich diversity, for example, by the types of benefits exchanged, the mechanisms of exchange, or the obligacy of each partner (reviewed in Bronstein 2015b, Douglas 2015).

Third, the development of population dynamic models for mutualism was stifled by the belief that simple mathematical approaches make unrealistic predictions. Foundational theory in community ecology developed from Lotka-Volterra models, which use linear functional responses to describe the effect of the interaction on each species. The Lotka-Volterra model can predict stable cycles (oscillations) for predator-prey interactions (Lotka 1925, Volterra 1926) or competitive exclusion for competition interactions (Volterra 1926, Gause 1934) – outcomes that stimulated fruitful empirical and theoretical work. In contrast, the Lotka-Volterra model for mutualistic interactions (Kostitzin 1934, Gause & Witt 1935) can predict unbounded population growth of both species (“the orgy of mutual benefaction,” May 1976). Theoretical work stagnated for nearly fifty years as authors attributed the supposed rarity of mutualism in natural ecosystems to the unstable nature of this outcome (e.g. Williamson 1972, May 1973, Goh 1979).

Population dynamic studies of mutualism began to reemerge in the 1980s, perhaps due to an increasing awareness of the prevalence and importance of mutualistic interactions (Boucher 1985, Bronstein 2015b). Authors showed that mutualism could be stabilized by incorporating negative density-dependence or mechanistic detail that explicitly limits benefit acquired from mutualism (see Table A1 of Appendix A). However, these models were criticized as either too case-specific to be useful or too abstract to be applicable (Bronstein 2015a). A handful of works have attempted to bridge this gap by organizing extant knowledge both conceptually and mathematically. For example, Addicott (1981) organized models by their effects on per-capita growth rate and/or equilibrium density for both participating species. Wolin and Lawlor (1984) categorized ways in which intraspecific density-dependence could limit benefits and stabilize mutualisms. Thompson *et al.* (2006) proposed a theoretical framework that organized mutualisms into those that affect birth rate, death rate, or habitat acquisition for each partner and predicted their ecological dynamics when immigration and emigration occur (i.e., in open systems). Most recently, Holland and DeAngelis (2010) categorized mutualisms as following “unidirectional” or “bidirectional” consumer-resource dynamics, in which one or both partners benefit from consuming costly resources provided by the other.

Despite these past advances, an “ecological theory of mutualism” has not penetrated into the greater ecological community (see recent textbooks, e.g., Gotelli 2008, Vandermeer & Goldberg 2013, Mittlebach & McGuill 2019). Calls continue for simple but usable theory that synthesizes among mutualisms to identify patterns in population dynamics and in the mechanisms that generate them (e.g. Addicott 1981, Callaway 2007, Bronstein 2015a). To that end, we develop theory in which the “benefits” of mutualism are an outcome of mechanisms that increase the per-capita growth rate of a population compared to that rate in the absence of its mutualistic partner. We derive models focusing on observable mechanisms so that each mathematical expression represents biological phenomena justified with empirical examples. Then, we synthesize a conceptual and mathematical framework that predicts population dynamics across mutualisms. Our work demonstrates that there is an ecological theory of mutualism that deserves attention from ecologists in general.

## Methods

Growth rate of any population *i* can be described as the difference between the population’s reproduction and mortality rates, which are functions of the density-independent per-capita birth (*b*_*i*_) and death (*d*_*i*_) rates, as well as per-capita self-limitation and other density-dependent processes (*s*_*i*_). In the absence of mutualism, we represent changes in population density (*N*_*i*_) over time (*t*) as:

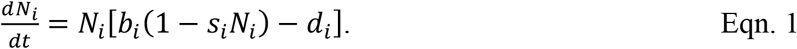

Eqn. 1 is continuous, deterministic, and ignores migration, which accommodates species with overlapping generations and allows us to focus on how the balance of births and deaths in a population leads to different dynamics, unobscured by stochasticity and the dynamics of other patches.

Nearly all mutualisms consist of an exchange of nutrition, protection, or transportation benefits. We derive and analyze models from Eqn. 1 for the most common exchanges: nutrition-for-nutrition, nutrition-for-protection, and nutrition-for-transportation (hereafter “nutrition,” “protection,” and “transport” mutualisms, respectively). Our equations for each type of benefit can be mixed-and-matched for different systems, like the protection-for-protection mutualism between clownfish and anemones (Table 1). Each of our models accommodates populations of both “obligate” mutualists that cannot persist in the absence of their partner and “facultative” mutualists that are self-sustaining. Table 2 summarizes our parameter definitions for all models.

**Table 1.** Population dynamic models for different types of mutualism. Mutualisms are grouped by the types of benefits exchanged (nutrition, protection, and transportation) between species (Sp. 1: ***N***_1_, Sp. 2: ***N***_2_), and their functional responses (M-M: Michaelis-Menten, H-II: Holling Type II). Additional models (with qualitatively similar results) are presented in Appendices A, C.

| Type of Mutualism | Examples<br>Sp. 1 / Sp. 2 | Dynamics of Sp. 1 | Dynamics of Sp. 2 |
| --- | --- | --- | --- |
| *Nutrition-for-Nutrition* | Plant / Microbe<br>(*mycorrhizae, rhizobia*) | Eqn. 2 w/ M-M | Eqn. 2 w/ M-M |
| *Nutrition-for-Protection* | Plant / Ant<br>Animal (*lycaenid caterpillars, scale insects, aphids*) / Ant | Eqn. 3 | Eqn. 2 w/ H-II |
| *Nutrition-for-Transportation* | Plant / Pollinator<br>Plant / Seed Disperser<br>(*ants, frugivores*) | Eqn. 4<br>Eqn. 5 | Eqn. 2 w/ H-II |
| *Protection-for-Protection* | Anemone / Clownfish | Eqn. 3 | Eqn. 3 |

**Table 2.**
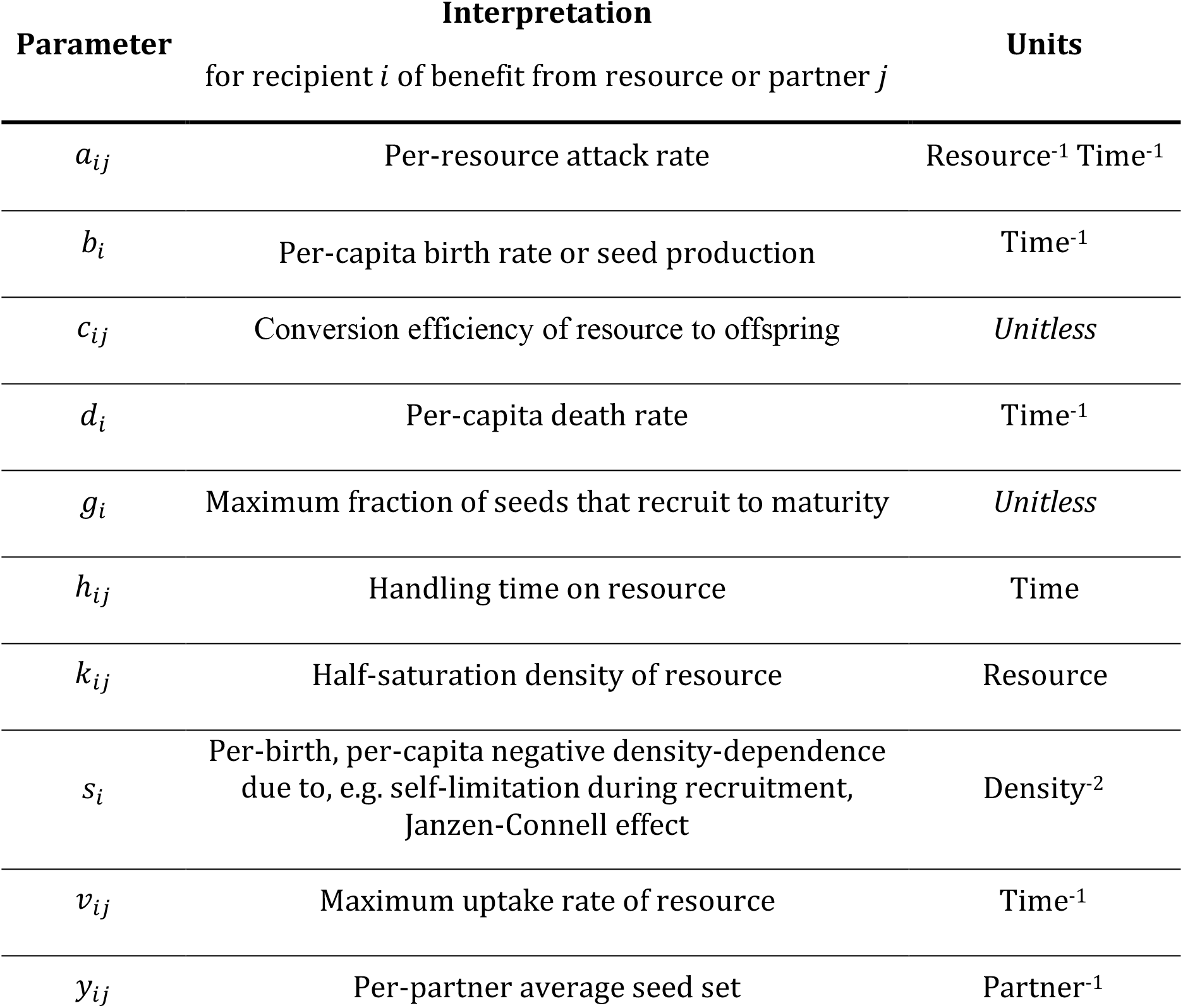
Table of paramteers. All parameters in our models (Eqns. 1–5 and Appendices) are assumed to be positive (> 0). The unit “Density” refers to population *i*, while the units “Resource” and “Partner” (depending on the system) refer to densities of population *j*. Subscripts *max*, *min*, and *diff* indicate the maximum, minimum, and difference, respectively, in a parameter value due to mutualism.

| Parameter | Interpretation<br>for recipient $i$ of benefit from resource or partner $j$ | Units |
| --- | --- | --- |
| $a_{ij}$ | Per-resource attack rate | Resource<sup>-1</sup> Time<sup>-1</sup> |
| $b_i$ | Per-capita birth rate or seed production | Time<sup>-1</sup> |
| $c_{ij}$ | Conversion efficiency of resource to offspring | Unitless |
| $d_i$ | Per-capita death rate | Time<sup>-1</sup> |
| $g_i$ | Maximum fraction of seeds that recruit to maturity | Unitless |
| $h_{ij}$ | Handling time on resource | Time |
| $k_{ij}$ | Half-saturation density of resource | Resource |
| $s_i$ | Per-birth, per-capita negative density-dependence due to, e.g. self-limitation during recruitment, Janzen-Connell effect | Density<sup>-2</sup> |
| $v_{ij}$ | Maximum uptake rate of resource | Time<sup>-1</sup> |
| $y_{ij}$ | Per-partner average seed set | Partner<sup>-1</sup> |

### Nutrition

A unifying feature of many mutualisms is that they involve nutritional mechanisms, in which the benefit to one species is gained by consuming nutritional rewards from its partner (Thompson 1982, Janzen 1985, Table 1 of Holland & DeAngelis 2010). For example, pollinators forage on nectar from flowers and mycorrhizal fungi uptake carbon from root nodules. We modify Eqn. 1 so that when *i* consumes the rewards of *j* with rate *C*_*R*_(***N***_*j*_), the per-capita growth rate of *i* increases proportionally:

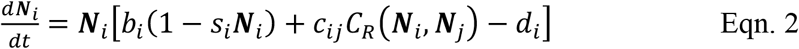

where *c*_*ij*_ is the conversion efficiency of *j*’s rewards to new individuals of *i*. If *i* forages for the rewards, the per-capita consumption 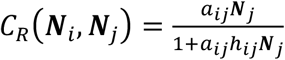, follows a Holling Type II functional response with attack rate *a*_*ij*_ and handling time *h*_*ij*_. If *i* uptakes the rewards directly (e.g., via diffusion), the per-capita consumption 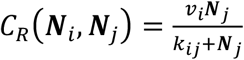 follows Michaelis-Menten kinetics with maximum uptake rate *v*_*ij*_ and half-saturation constant *k*_*ij*_ (i.e., the density of *j* at which half the maximum uptake rate is achieved). Both forms encode the reasonable assumption that *i*’s consumption rate on *j*’s rewards saturates with increasing density of *j*. Indeed, the expressions are identical when *a*_*ij*_ = *v*_*i*_/*k*_*ij*_, *h*_*ij*_ = 1/*v*_*i*_. Species *i* is an obligate partner of *j* when *r*_*i*_ = *b*_*i*_ − *d*_*i*_ < 0; otherwise (*r*_*i*_ ≥ 0), *i* is facultative.

Eqn. 2 is the most similar of our models to previous theory. It is an extension of the consumer equation of Rosenzweig and MacArthur’s (1963) model when *b*_*i*_ > 0, and identical to Wright’s (1989) model in which nutritional benefits saturate as a result of constraints on handling time during foraging. It is also identical to Holland and DeAngelis’ (2010) equation for consumer mutualists that do not supply rewards (Table A1).

### Protection

Protection mechanisms characterize a wide array of interactions in which the presence or behavior of a species protects another from natural enemies. The best-studied protection mutualisms are between ants and rewards-provisioning species like plants, lycaenid caterpillars, scale insects, and aphids (Ness *et al.* 2010). Ants harvest rewards like nectar, food bodies, and honeydew while cultivating and defending their resources through deterrence, attendance, and direct attack. Though these protection services have been observed to increase reproduction, the primary benefit to the resource species is a reduction in mortality due to, e.g., natural enemies including herbivores, predators, and parasitoids (Ness *et al.* 2010, Trager *et al.* 2010). Therefore, we assume that benefits of protection services (*S*) are exclusively reductions in per-capita death rate. We model these reductions as the difference between the maximum per-capita death rate of *i* inflicted by its natural enemies 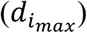 and the protection services provided by *j*, 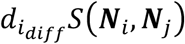, which incorporated in Eqn. 1 yields:

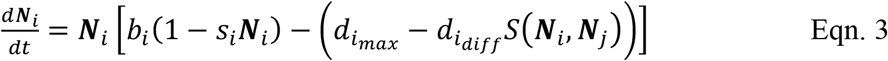

where 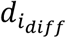 is the maximum reduction in *i*’s death rate caused by *j*’s protection, with minimum death rate 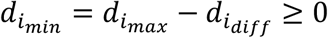. Species *i* is obligate upon *j* when *r*_*i*_ = *b*_*i*_ − *d*_*i*_ < 0; otherwise *i* is facultative.

We assume that the protector population *j* benefits from foraging on rewards provided by *i* and, therefore, model its dynamics with Eqn. 2 using the foraging functional response (Table 1). If *j* reduces mortality via simple deterrence (“scaring off” natural enemies), we choose 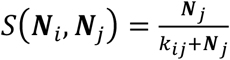 so that per-capita death rate declines proportional to *j*’s perceived abundance around the recipient species, saturating when *j*’s density is high. We consider other mechanisms of protection in Appendix C.

The nutrition (Eqn. 2) and protection (Eqn. 3) equations are identical when using the Michaelis-Menten functional response, with 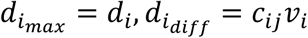. Eqn. 3 is also similar to Thompson *et al.*’s (2006) model for protection mutualisms in open systems, when immigration and habitat provisioning are assumed to be negligible (Table A1).

### Transport

Transportation mechanisms characterize the best studied mutualisms, perhaps due to the astonishing diversity by which immobile species attract more-mobile species to disperse their gametes. In the most common interactions (pollination and seed dispersal), animals visit plants to feed on rewards (including nectar and fruits), providing reproductive services (transport of pollen and seeds) incidentally during foraging. Except in special cases (e.g., male bees visiting flowers to collect “perfume”), these rewards offer primarily nutritional benefit (Willmer 2011). Thus, we model the dynamics of transporter populations (*i*) using Eqn. 2 with the foraging functional response (Table 1).

For plant populations (*i*) that receive reproductive services (*S*), we define *b*_*i*_ as per-capita seed set, *g*_*i*_(> 0, ≤ 1) as the constant fraction of total seed set that germinates, and *s*_*i*_ as negative density-dependence (e.g., Janzen-Connell effect, seed competition for recruitment). Plant *i* is obligate upon *j* when *r*_*i*_ = *g*_*i*_*b*_*i*_ − *d*_*i*_ < 0; otherwise *i* is facultative.

Reproductive services are functions of *j*’s visitation to *i* (see below) and increase the number of mature individuals in *i*. We derive separate models for pollination services (which increase seed set) and seed dispersal services (which decrease mortality during recruitment). We assume that foraging rate on rewards (*C*_*R*_) is a good proxy for visitation rate. Though some visits are unsuccessful due to depleted rewards or deception, we consider this mathematically similar to a predator unsuccessfully attacking a prey.

Modifying Eqn. 1 for pollination benefits yields:

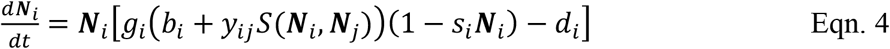

where *y*_*ij*_ is the conversion efficiency of pollination services to seed production. Pollination requires the transfer of pollen from one individual’s flower to the stigma of a conspecific plant individual, which occurs when animals visit flowers to forage. We therefore assume 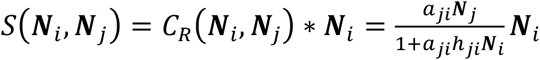 where the contribution of the animal population (*j*) to per-capita seed set is proportional to *j*’s total foraging rate on *i*, calculated by multiplying *j*’s per-capita foraging rate on *i*’s rewards with plant density (***N***_*i*_). This expression accounts for the repeated interactions between plant and animal individuals required for conspecific pollen transfer (Vázquez *et al.* 2005).

In seed dispersal interactions, animals visit plants to forage on fruit or elaiosomes, later depositing the seed away from the parent plant. This process may increase germination rate by improving seed condition during passage through dispersers’ guts, but more commonly increases recruitment by lessening density-dependent sources of mortality due to predators and pathogens, which are most abundant near adult plants (Fricke *et al.* 2013). We assume that negative density-dependence 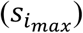 can be reduced to a minimum of 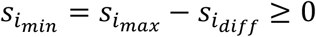 when dispersal services (*S*) are maximally effective. Modifying Eqn. 1 yields:

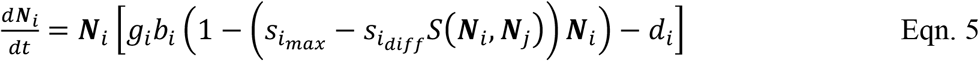

We assume 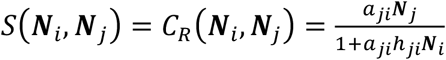, where the contribution of animals (*j*) to per-capita seed set is proportional to *j*’s per-capita (instead of total) foraging rate on *i*, because repeated interactions between plant and animal individuals are possible but not required for effective seed dispersal (as opposed to the requirement for effective pollination). To ensure that 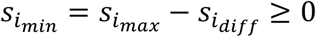, we choose parameter values that fulfill 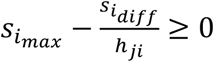

Eqn. 5 differs from previous models mathematically, but is conceptually similar to earlier theory that modeled mutualistic benefits as increasing carrying capacity (Whittaker 1975, Dean 1983, Wolin & Lawlor 1984, Neuhauser & Fargione 2004, Graves *et al.* 2006, see Table A1), which is analogous to reducing the magnitude of negative density-dependence. However, our pollination model (Eqn. 4) differs substantially from previous work in the two-species population dynamic literature (Table A1). Its uniqueness is due to the conceptually-important choice for negative density-dependence to limit (using a multiplied term) the fraction of fertilized ovules that survive as seedlings to mature to reproductive adults. If we instead assume that negative density-dependence limits overall per-capita growth (using an additive term) as in previous works, our pollination and seed dispersal models would be mathematically identical.

### Rewards and Other Costs of Mutualism

Species that receive services often offer nutritional “rewards” to attract their partner. These rewards are often partitioned from other components of the individual and have a unique chemical composition. Plants can produce nectar, a simple sugary solution, in flowers or nectaries separate from their vegetation, which is constructed from more complex carbohydrates and defensive compounds. Aphids and other hemipterans secrete honeydew that can be harvested separately from the rest of their body mass without killing individuals in the population. Nutritional mutualisms also follow this rewards paradigm with species provisioning a non-limiting resource in exchange for the acquisition of a limiting resource (Bronstein 2009).

Are rewards costly to offer? Evidence is sparse. Some rewards are waste products (Ness *et al.* 2010, e.g. honeydew is partially-digested phloem) that can even harm individuals if not removed (Gullan 1997), while other rewards are costly to individuals to construct (Brandenburg *et al.* 2012) or to have exploited (Yao & Akimoto 2001). However, these individual-level costs are likely to be buffered at the population level (see discussion in Appendix A). We therefore assume that costs associated with rewards do not substantially impact population density or, if they do, the impact could be represented as small changes to per-capita birth and death rates approximated as constant with respect to the producing species’ density. Other costs including increased handling time when foraging or increased exposure to parasites transported by visitors are also likely to be negligible or fixed at the population level (i.e., absorbed in parameter values, Holland & DeAngelis 2010). Therefore, we do not include explicit cost terms in our models.

Nonetheless, individual-level costs that scale with rewards exploitation are often assumed to have population-level impacts (e.g. Aizen *et al.* 2014). To understand the impact of this assumption, we compare the dynamics of our simpler models to extended models with rewards exploitation (see Appendix C and discussion of overexploitation dynamics, below).

## Results

We found that mutualisms have important similarities in their dynamics and stability. First, our models always predict stable coexistence when both species are facultative (Fig. 1F, 2F-G, 3F-G). Second, threshold or Allee effects may destabilize the system when at least one species is obligate and one or both partners are at low density (Figs. 1–3, left-column panels). Then, depending on initial conditions, both species go extinct or the facultative species persists at low density in the absence of its partner. Third, our models always display threshold effects when both partners are obligate which result in extinction at low densities (Fig. 1A, 2A, 3A).

**Figure 1.**
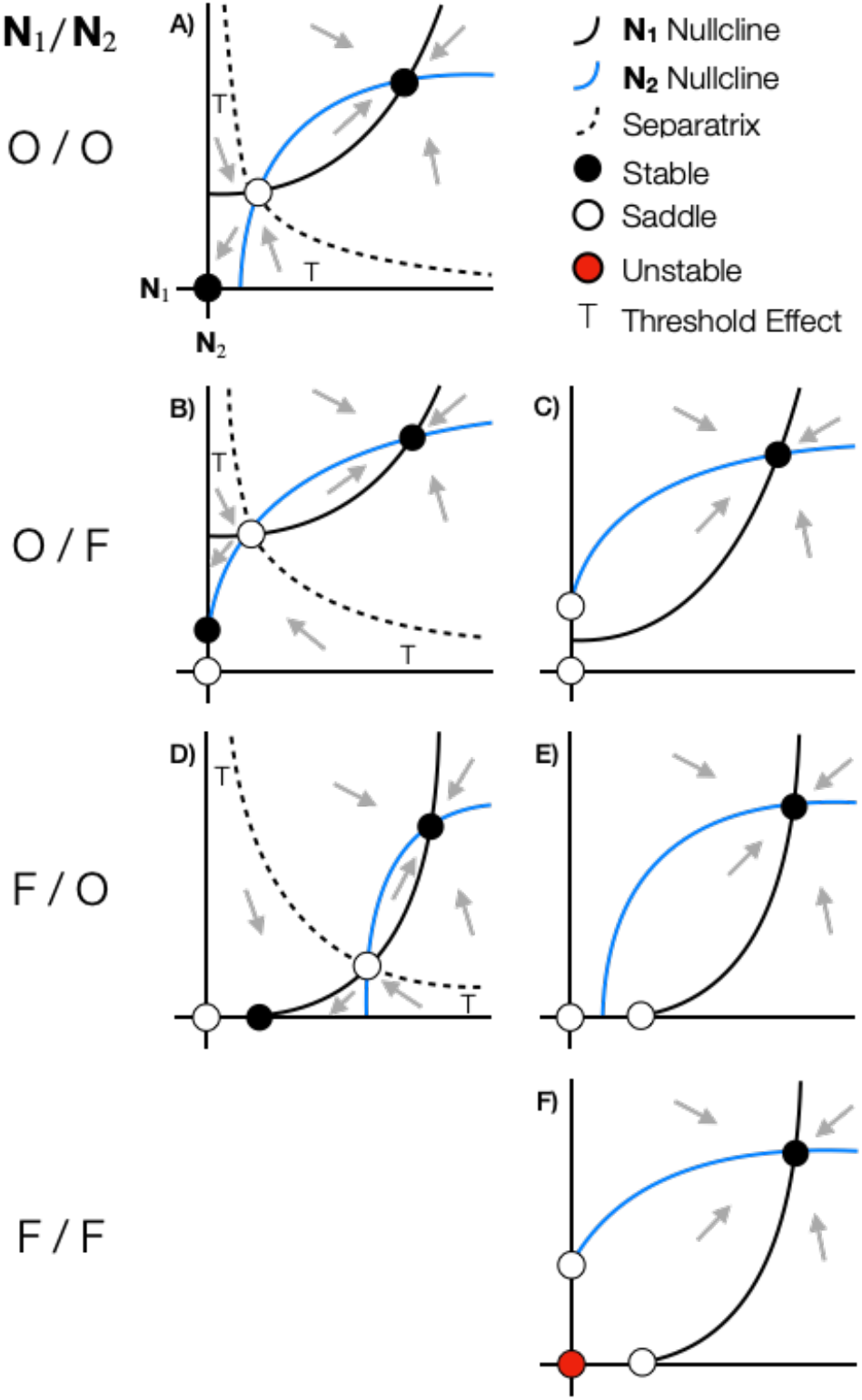
Phase plane diagrams for nutritional and protection mutualisms. Curves represent nullclines along which population growth for each species is zero. Trivial nullclines (at ***N***_1_ = 0, ***N***_2_ = 0) are omitted for clarity. Rows illustrate all possible qualitative dynamics when coexistence is feasible for each combination of obligate (O) and facultative (F) partners. The S and Q axes are the densities of species 1 (***N***_1_) and 2 (***N***_2_), respectively. Only the quadrant with positive abundances is shown. Arrows illustrate the direction of population change in each region of the phase plane. Equilibria occur where the nullclines for ***N***_1_ (black) and ***N***_2_ (gray) intersect. Filled black equilibria are stable attractors, filled red equilibria are unstable repellers, and hollow equilibria are half-stable saddle points. Here, saddle points induce *threshold effects* (T) in which one population declines under a threshold of its partner’s density (the separatrix, dashed line), leading to *single-species persistence* of the facultative partner (**B**, **D**) or *extinction* when both partners are obligate (**A**). Above the threshold (in **A**, **B**, **D**) or in systems without a threshold (**C**, **E**, **F**), the system always achieves *stable coexistence*. In fact, stable coexistence is the only possible outcome when both partners are facultative (**F**) or in some configurations of nullclines when one partner is facultative and the other obligate (**C**, **D**).

**Figure 2.**
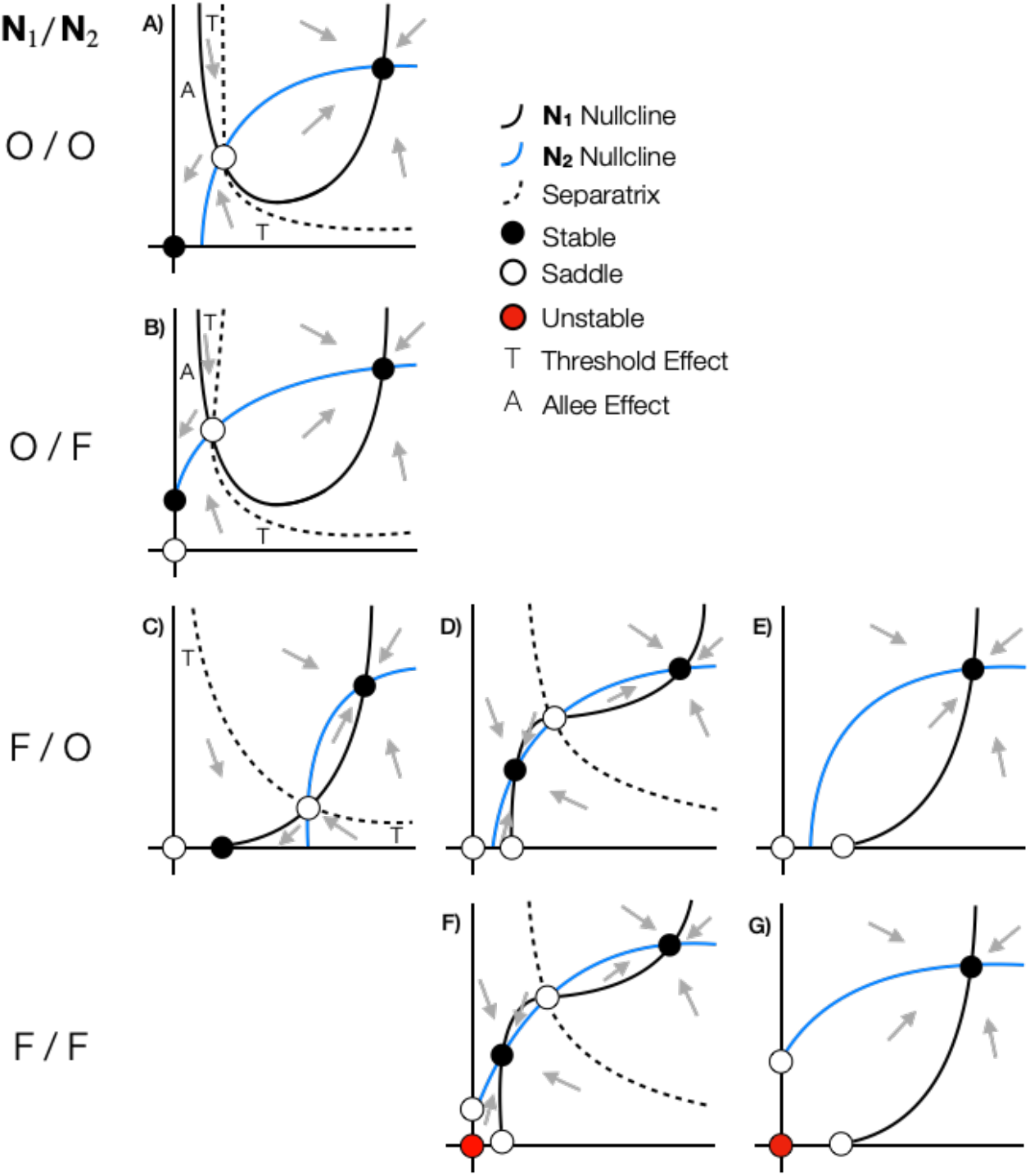
Phase plane diagrams for pollination mutualisms. All formatting and terminology follows Fig. 1. Here, saddle points can induce threshold effects (T) when at least one species is obligate (**A, B, C**). When species’ densities are above the threshold, they achieve stable coexistence; below the threshold, the system collapses to extinction (**A**) or single-species persistence (**B, C**) of the facultative partner. When the plant partner (***N***_1_, black) is obligate (**A, B**), threshold effects additionally lead to strong, demographic Allee effects (A, Kramer et al. 2009) in which the plant population declines under a threshold of its own density. Saddle points can also induce bistability (i.e., two stable coexistence equilibria, **D, F**), where species stably coexitthe plant nullclines betweens at the low-density equilibrium if their density were initially below the separatrix (dashed line) or at the high-density equilibrium otherwise. Finally, when plants are facultative, stable coexistence is the only possible outcome for some nullclines configurations (**E**, **G**).

**Figure 3.**
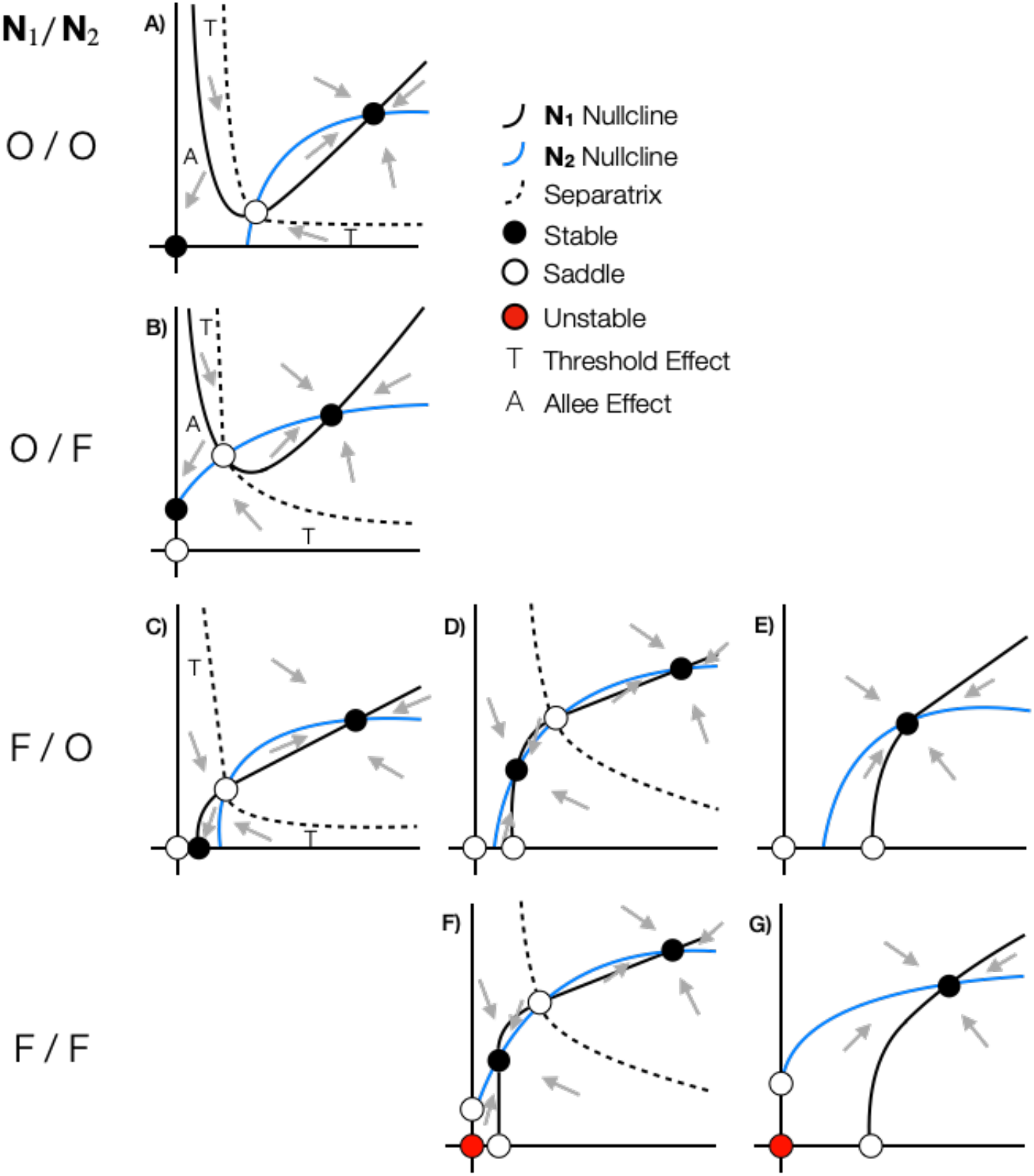
Phase plane diagrams for seed dispersal mutualisms. All formatting and terminology follows Figs. 1–2. All dynamics described in Fig. 1 also apply here. Pollination and seed dispersal mutualisms differ only the geometry of the plant nullcline (***N***_1_, black), but not in their dynamical outcomes.

Below we give a detailed description of the dynamic outcomes of each of our models, assuming coexistence is feasible. We organize our results by common and unique outcomes including Extinction, Single-Species Persistence, Coexistence, Threshold Effects, Allee Effects, and Bistability (see definitions below and Appendix B for mathematical details).

### Nutrition and Protection

Nutrition and protection mutualisms have the same mathematical form and identical dynamical results despite their different derivations. We call ***N***_1_ the plant (or attendee) population and ***N***_2_ the microbe (or protector) population for nutrition (or protection) mutualisms (Table 1). Microbial and protector nullclines (gray curves, Fig. 1) are concave down, increasing functions that saturate with respect to plant or attendee density. The plant and attendee populations follow the same dynamics yielding symmetrical nullclines (black curves, Fig. 1). Together, these result in the following outcomes:

1. **Extinction** (Fig. 1, all panels, circles at the origin). The extinction equilibrium is attracting (a stable point) only when both partners are obligate (Fig. 1A). Otherwise, one or both populations are repelled from extinction. The origin is a saddle point for facultative-obligate pairs (Fig. 1B-E) or an unstable point when both partners are facultative (Fig. 1F).
2. **Single-Species Persistence** (Fig. 1B-F, circles on axes). Facultative species can persist at density *r*_*i*_/*b*_*i*_*s*_*i*_ in the absence of their partner. These single-species equilibria are attracting when threshold effects are present (Fig. 1B, 1D). Otherwise (Fig. 1C, 1E, 1F), they are saddle points and the system will tend towards stable coexistence.
3. **Stable Coexistence** (Fig. 1, all panels, off-axes black circles). The coexistence equilibrium is a stable node (no oscillations) at higher density than either species could achieve alone. Increased density past this equilibrium in either species causes its population to decrease due to negative density-dependence. Therefore, the “orgy of mutual benefaction” (May 1976) cannot occur.
4. **Threshold Effects** (Fig. 1, left-column panels, “T”). The nullclines intersect twice under some parameterizations when at least one species is an obligate partner. This yields a saddle point (Fig. 1A, 1B, 1D, off-axes unfilled circle) at densities intermediate to the stable coexistence and single-species persistence or extinction equilibria. The saddle determines a threshold (the “separatrix,” dashed line) below which obligate partners go extinct even if initially highly abundant (e.g., left “T” region in Fig. 1d). This means that one species is too low in density to provide sufficient benefits to its partner, causing the partner’s population to decline. The low-density species continues to benefit from mutualism but its increase in density cannot occur fast enough to save the system from extinction. Above the separatrix, one or both species are of high enough density that benefits from mutualism cause positive population growth in their partners and the system will achieve stable coexistence. Alternative protection mechanisms of different mathematical forms yield qualitatively similar results (Appendix C).

### Transport

We call ***N***_1_ the plant population and ***N***_2_ the animal population (Table 1). The animal nullclines (gray curves) are identical to the microbe/protector nullclines in Fig. 1. The plant nullclines in the pollination and seed dispersal models are both top-open humps bounded on the left side by a vertical asymptote at zero (Figs. 2A-B, 3A-B) despite different derivations. These nullclines differ in their high-density behaviors, where the pollination nullcline is concave up and saturates to 1/*s*_1_ (Fig. 2), while the seed-dispersal nullcline is concave down when facultative (Fig. 3C-G) and continues to increase with a linear slope 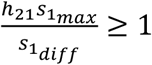

Differences in the plant nullclines between seed dispersal and pollination interactions manifest in different nullcline geometries but similar dynamical outcomes. Many of the outcomes are similar to those described for nutrition and protection mutualisms, which we do not restate. We describe only dynamics unique to the transport models:

1. **Extinction** (Figs. 2–3, all panels, circles at the origin).
2. **Single-Species Persistence** (Figs. 2B-G, 3B-G, circles on axes). Facultative plants can persist at density *r*_1_/*g*_1_*b*_1_*s*_1_ in the absence of their partner.
3. **Stable Coexistence** (Figs. 2–3, all panels, off-axes black circles). One stable coexistence equilibrium is present when plants are obligate mutualists (Fig. 2A-B, Fig. 3A-B). Two stable coexistence equilibria (bistability) are possible when plants are facultative (Fig. 2D, 2F, 3D, 3F).
4. **Bistability** (Figs. 2–3, middle-column panels). When plants are facultative, the plant and animal nullclines intersect three times under some parameterizations. This yields a saddle point (off-axes unfilled circle) bisected by a separatrix (dashed line) that divides the plane into two basins of attraction corresponding to a lower- and higher-density stable coexistence, both at higher density than either species could achieve alone. The system will tend to either coexistence equilibrium depending on initial conditions.
5. **Threshold Effects** (Figs. 2–3, left-column panels, “T”).
6. **Allee Effects** (Figs. 2A-B, 3A-B, “A”). When plants are obligate, their population declines under a threshold of their own density regardless of the density of their partner. This results from the asymptotic behavior of the plant nullcline at ***N***_1_ = 0, which occurs by different mechanisms depending on the transport model. In the pollination model, it occurs because benefit is proportional to the total consumption rate by animals, whereas in the seed-dispersal model it occurs because benefit affects the density-dependent term.

## Discussion

Previous models have been criticized as too case-specific or too phenomenological to further empirical and theoretical understanding of generalities in the ecologies of diverse mutualisms (Bronstein 2015a). To address this, we derived models that balance mechanistic detail with the potential for mechanisms to have population-level effects. We found that predictions for the population dynamics of mutualisms are qualitatively robust across derivations, including level of detail, types of benefit, and inspiring systems. We now organize these predictions in a (novel) theoretical framework that synthesizes our conceptual and mathematical results (Table 3, Fig. 4). We then highlight the assumptions that restrict the applicability of our work and identify how these assumptions can be relaxed. We conclude that a coherent and well-developed ecological theory of mutualism exists.

**Table 3.** Theoretical framework. Synthesis of Conceptual and Mathematical results for each type of mutualism. All entries are with respect to the recipient of mutualistic benefits (Eqns. 2–5) and assume that the partner is a nutritional mutualist as in our Figs. 1–3. Dashes (−) indicate no notable behavior in that region. The mapping of different “Types of Benefits” to “Processes” and “Mechanisms” is a conceptual simplification that represents general patterns among mutualisms. Mechanisms can result from different behaviors of partners and recipients; we specify one characteristic behavior for each of our models (“Means of Effect”). In all cases, per-capita benefits are a function of partner density. Additionally, the “Effects” of mutualism are either directly dependent or independent of recipient density. All effects are saturating, either due to interspecific density-dependence (nutrition, protection) or intraspecific density-dependence (transport, alternate protection models [Appendix C]). These distinctions, in addition to whether the recipient species is obligate (O) or facultative (F), determine different “Nullcline Geometries” and “Dynamical Outcomes”. For example, both models with “density-dependent” effects have nullclines with a vertical asymptote at zero when obligate, and can change concavity between low and high density when facultative. Different dynamical outcomes are dependent on parameterization of the model, the type of partner species, and whether the partner is obligate or facultative. Outcomes are separated into columns (headed O/F) when dependent on partner obligacy (highlighted with a superscripted *P*) and are separated by line-breaks when dependent on parameterization. In all our models, stable coexistence can always occur at higher density than either species could achieve alone. Our models are thus our distinguished by their low density-behavior: “T” and “A” refer to threshold and Allee effects, respectively. “B” refers to stable coexistence at low density, which gives two stable coexistence equilibria (bistability). When facultative, species can also potentially persist at a low-density, single-species equilibrium (not included in the table).

| CONCEPTUAL |  |  |  |  | MATHEMATICAL |  |  |  |  |  |  |
| --- | --- | --- | --- | --- | --- | --- | --- | --- | --- | --- | --- |
| Type of Benefit | Process | Mechanism | Means of Effect | Effect | Nullcline Geometry |  |  | Dynamical Outcomes |  |  |  |
|  |  |  |  |  | Obligacy (recipient) | Low Density | High Density | Low Density |  | High Density |  |
|  |  |  |  |  |  |  |  | O<sup>P</sup> | F<sup>P</sup> | O<sup>P</sup> | F<sup>P</sup> |
| *Nutrition* | Consumption | Consumption | **Per-capita** consumption **on** partner | Density-independent (↑ *b<sub>i</sub>*, ↓ *d<sub>i</sub>* or both) | O | – | Saturating, concave up | T | T | Stable coexistence |  |
|  |  |  |  |  | F | – |  | T | – |  |  |
| *Protection* | Mortality | ↓ Mortality | **Per-capita** deterrence/attendance **by** partner | Density-independent (↓ *d<sub>i</sub>*) | O | – | Saturating, concave up | T | T | Stable coexistence |  |
|  |  |  |  |  | F | – |  | T | – |  |  |
| *Transportation:*<br>Pollination | Reproduction | ↑ Seed production | **Total** consumption **by** partner | Density-dependent (↑ *b<sub>i</sub>*) | O | Asymptote at 0 | Saturating, concave up | T, A | T, A | Stable coexistence |  |
|  |  |  |  |  | F | Changing concavity |  | T, A<br>B | B |  |  |
| Seed dispersal | Reproduction | ↓ Density-dependent mortality during recruitment | **Per-capita** consumption **by** partner | Density-dependent (↓ *s<sub>i</sub>*) | O | Asymptote at 0 | Linear growth | T, A | T, A | Stable coexistence |  |
|  |  |  |  |  | F | Concave down |  | T, A<br>B | B; |  |  |

**Figure 4.**
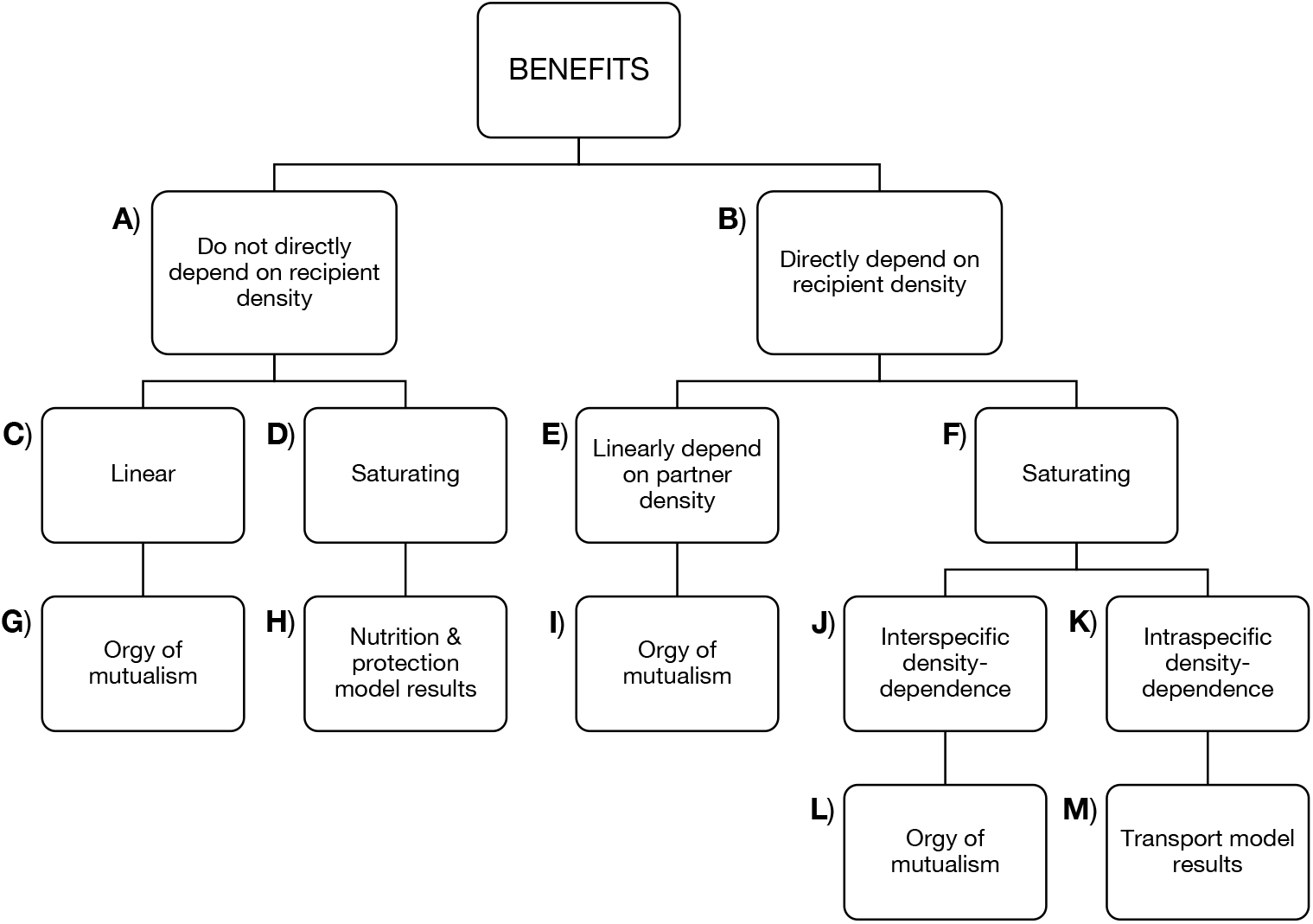
An ecological theory of mutualism. By synthesizing the results of our models with historical models derived with varying levels of mechanism for diverse motivating systems (Appendix A), we demonstrate that ecological models of mutualism have predictable dynamical outcomes. This synthesis assumes that benefits of mutualisms strictly increase per-capita growth rate of the recipient population, affect only a single ecological process, and depend on partner density. Benefits may furthermore (**B**) directly depend or (**A**) not directly depend on recipient density. (**C**) If benefits simply accumulate linearly as function of partner density (i.e., do not directly depend on recipient density), models predict (**G**) unstable coexistence – the “orgy of mutual[ism].” (**D**) If benefits instead saturate (either due to interspecific density-dependence, intraspecific density-dependence, or both), mutualisms exhibit (**H**) the same dynamical outcomes as our nutrition and protection models. That is, stable coexistence and threshold effects when one partner is obligate. Many historical models fall into this category and predict identical outcomes. (**B**) When benefits directly depend on recipient density, they may additionally (**E**) accumulate linearly as a function of partner density, leading to (**I**) the orgy of mutualism, or (**F**) saturate. (**J**) If benefits saturate with respect to partner density, models predict (**L**) the orgy of mutualism. (**K**) If benefits instead saturate with respect to recipient density, as in our pollination and seed dispersal models, mutualisms exhibit (**M**) stable coexistence, threshold effects when one partner is obligate, Allee effects when plants are obligate, and potential bistability when plants are facultative. The combination of both direct dependence and saturation in terms of recipient density in our transport models appears to be unique among historical models. See Appendix A for historical models and descriptions of their dynamical outcomes.

### Synthesizing Mechanisms in a Mathematical Framework

We followed previous conceptual work to develop models for species benefiting from nutritional, protection, and transportation mechanisms, as these putatively mapped onto the processes of consumption, mortality, and reproduction (Table 3). Our mathematical results revealed that nutritional and protection mechanisms are similar to each other because they modify density-independent rates (per-capita birth, death, or both) by saturating processes directly dependent on partner density or per-capita behavior rates (i.e., partner or recipient consumption, deterrence, attendance; see Appendices A, C). Similarly, transportation mechanisms are similar to each other because they both directly depend on and saturate in terms of recipient density, either because they are proportional to the partner’s total behavior rates (pollination) or because the partner’s per-capita behavior modifies a density-dependent rate (seed dispersal).

This suggests a more informative way to group mutualisms: by whether their effects are directly dependent on recipient populations (Fig. 4). This scheme predicts that some interactions will be dynamically similar despite differing in their types of benefit. For example, protection services in some systems may reduce density-dependent sources of mortality such as disease (e.g., cleaning by ants reduces aphid mortality, Nielsen *et al.* 2010, Durak *et al.* 2016). Modeling this similarly to seed dispersal, where benefits are directly dependent on recipient density, predicts an Allee effect at low density in obligate partners (Appendix C). Alternatively, when its effects are not directly dependent on recipient density, mutualism will have indistinguishable impacts on population dynamics, whether resulting from nutritional, protection, or transportation mechanisms (of the same functional form). For example, seed dispersal services (per-capita visitation) may increase (density-independent) germination rate (*g*_*i*_) in some systems (e.g., Fricke *et al.* 2013). Modeling this yields qualitatively identical results to our nutrition and protection models (Appendix C).

### Synthesizing Population Dynamic Behavior

All four mutualism models studied here exhibit similar dynamical outcomes (Table 3), which were also predicted by several historical models (Appendix A). Specifically, when feasible, coexistence is stable, non-oscillatory, and populations grow with bound. All obligate-obligate mutualisms and certain interactions between obligate and facultative mutualists exhibit thresholds, under which the low density of one partner destabilizes the system.

Mutualisms are commonly modeled as having effects independent of recipient density (Appendix A). We modeled the benefits provided by pollination and seed dispersal as both directly dependent on and saturating in terms of recipient (plant) density, yielding the additional dynamical outcomes of Allee effects or bistability when plants are obligate or facultative, respectively. Other mutualisms may exhibit these outcomes if they: i) modify a density-dependent process, ii) affect a total rate (e.g., visitation) on the recipient population, or iii) include a facultative partner that follows a Type III functional response (Appendix C).

Understanding how frequently the effects of mutualisms are directly dependent or independent of recipient density is an important avenue for future work. For example, empirical work could distinguish between these effects by testing for a dynamical transition from threshold to Allee effects in obligate plants when their pollinators are at very high density (compare Fig. 1A-B to Fig. 2A-B).

Our nutrition and protection models are identical to those proposed by Holland and DeAngelis (2010) when excluding their cost terms for supplying resources (like rewards). The cost terms in their models lead to lobe-shaped nullclines and substantially different dynamical outcomes. Specifically, our models (no cost terms) predict a threshold effect where the high-density partner decreases and the low-density partner increases, while their models (with cost terms) predict overexploitation where the high-density partner increases and the low-density partner decreases (shifting the interaction from mutualism to parasitism), eventually causing an Allee effect and extinction of one or both partners. Empirical studies could test these alternative predictions by tracking population sizes over time when one partner is at low density.

Interestingly, overexploitation in their models occurs due to the increased costs to the exploited partner of supplying resources. Extensions to our models with exploitation (as opposed to rewards production) costs can also predict overexploitation dynamics with fewer parameters (Appendices C).

### Population-Level Impacts of Benefits and Costs Occurring at the Individual Level

Population dynamic models of mutualistic interactions, including ours, assume that mutualisms have population-level impacts. However, most empirical studies quantify the benefits and costs of mutualisms at the individual level in terms of fitness or even by using a single proxy for fitness (Bronstein 2001, Ford *et al.* 2015). Those effects do not necessarily imply population- and community-level impacts of mutualism (Williamson 1972, Flatt & Weisser 2000, Ford *et al.* 2015, Fredrickson 2015). Therefore, empirical work on population dynamics of mutualisms is of foremost importance to evaluate historical and current ecological theory on mutualisms.

To derive our models, we synthesized available knowledge across mutualisms on population-level costs of offering rewards (Appendix A). Using this synthesis, we assumed that costs associated with rewards did not have population-level effects or that they were of low or “fixed” cost which could be accounted for in demographic parameters. In contrast, high magnitude or “variable” costs that are a function of partner density should be accounted for explicitly (Holland et al. 2002, Morris *et al.* 2010). More generally, we expect that “construction” costs (attracting partners, rewards production) are more likely to be negligible than “exploitation” costs (rewards consumption, interference, destruction, mortality; Holland *et al.* 2002) which scale with a partner’s visitation rate, and thus are more likely to have impacts on the recipient population (see extensions, Appendix C). Similar conceptual and empirical syntheses among mutualisms are needed to provide insight into how mechanisms that affect fitness also impact population dynamics.

### Applying Our Models

Our simple models can be applied to a variety of mutualisms. This final subsection discusses our most important assumptions in order to clarify the conditions under which our models apply. See Gotelli (2008) for a review of basic assumptions of population dynamic models and their consequences.

#### Benefit via a single mechanism

The predictions of our models apply when there is one primary mechanism of mutualistic benefit to each partner. For example, we assume that a nutritional mutualist benefits exclusively from consuming its partner’s rewards – not other components of the partner’s biomass and not from other aspects of the interaction that affect different vital rates. An interesting extension of our work would be to synthesize population dynamic behavior for “double mutualisms,” in which there are two mechanisms of mutualistic benefit to a partner (Fuster *et al.* 2019). Double mutualisms may play important ecological roles in ecosystems with limited resources, such as islands.

#### Negative density-dependence

We made the broad assumption that resources for reproduction and growth are limited, so vital rates are dependent on population size (Gotelli 2008). We included a negative density-dependence term in each of our population dynamic equations to represent, e.g., self-limitation due to competition or the Janzen-Connell effect (a common assumption, Table A1). The combination of negative density-dependence and saturating functional responses in our models curves the nullclines so that they intersect and result in stable coexistence.

#### Functional forms

Density-dependent functional responses limit positive feedbacks that cause unbounded population growth (the “orgy of mutual[ism]”) observed, e.g., in Lotka-Volterra models with density-independent (linear) functional responses. Historically, authors focused on intraspecific density dependence, where per-capita benefits decrease and saturate with increasing recipient density (but see Appendix A). Recent work advocates for interspecific density dependence, where per-capita benefits saturate with increasing *partner* density, to integrate mutualism into broader consumer-resource theory (Wright 1989, Holland & DeAngelis 2010, Holland 2015). Our work primarily studied saturating (Holling Type II and Michaelis-Menten) functional responses with either inter- or intraspecific density-dependence (also see Appendix C). We used interspecific density-dependence for nutritional and protection mutualists, where recipients benefit from consuming their partners’ rewards or from deterrence of enemies by their partners, respectively. We used intraspecific density-dependence for transportation mutualists, where recipients benefit from their partners’ consumption of their own rewards. However, the distinction between intra- and inter-specific density dependence is not predictive of different population dynamics when the effects of mutualism do not directly depend on recipient density (Fig. 4). For example, protection services could instead be modeled with intraspecific density dependence if benefits are proportional to protectors’ recruitment on recipients (e.g., Morales 2000). This does not modify the qualitative dynamics of our protection model (Appendix C).

In summary, we proposed a theoretical framework that ties mechanisms to predicted population dynamics (Table 3, Fig. 4). The predictions of this framework apply when population-level benefits accumulate via a single mechanism and when population-level costs are negligible, are fixed effects, or simply diminish population-level benefits. Additionally, functional forms should be chosen according to the system. Then, effects of mutualism could potentially be identified experimentally by testing where per-capita benefits increase proportionally to or are independent of recipient density, respectively. Finally, our models could be falsified by assessing transient population dynamics for threshold effects, Allee effects, or overexploitation when one species is at low density.

## Conclusion

A critical part of elevating mutualism to the “third pillar of ecology” is developing theory that underpins fitness, population dynamics, and community dynamics of mutualistic interactions – but work on population dynamics is notably underrepresented. Here, we united conceptual mathematical approaches to identify patterns in the processes that generate mutualisms and the population dynamics that result. Despite different derivations, mechanisms, and inspiring systems, our models and many historical models make similar qualitative predictions when grouped by whether the effects of mutualism are directly dependent or independent of recipient density. This suggests that there exists a robust population dynamic theory of mutualism that can make general predictions. These predictions (including stable coexistence, threshold effects, Allee effects, and overexploitation) can be tested by combining empirical and theoretical approaches (e.g. Breton & Addicott 1992, Morales 2000, Palmer *et al.* 2010, Kang *et al.* 2011, Ford *et al.* 2015, also see Holland 2015). Such work will contribute to our understanding of general patterns and processes within and among mutualisms, and inform efforts to preserve mutualistic systems (Bronstein 2001, Callaway 2007, Bronstein 2015a,b).

## Acknowledgements

This research was supported by National Science Foundation Graduate Research Fellowship DGE-1143953 to K.R.S.H., National Science Foundation Graduate Research Fellowship DGE-1256260 to D.P.M., and National Science Foundation grant DEB-1834497 to F.S.V.

## Appendix A Additional Discussion

## Costs of Rewards at the Population Level

Offering rewards to mutualistic partners can incur costs to individuals (Bronstein 2001). To our knowledge, however, there are not empirical studies that have measured how these costs result in changes to population growth. Given this lack of empirical information, we chose to assume that costs associated with rewards are negligible at the population level (or can otherwise be accounted for in parameter values). This is justified for at least four reasons: First, costs associated with rewards can simply act to reduce benefit provided by mutualism (e.g., reduced seed set due to nectar production, Brandenburg *et al.* 2012). Second, rewards and the organs that produce them are typically ephemeral in comparison to individuals in the population. They can be damaged or decay, but, instantaneously, these costs are unlikely to result in death or damage to individuals (Revilla 2015). Third, many costs associated with rewards are “fixed” in that they are independent of partner density (e.g., sunk costs of molecular machinery used to produce rewards, Holland *et al.* 2002) or are otherwise compensated for within individuals (e.g., carbon costs of supporting microbial mutualists stimulate compensatory photosynthesis, Kaschuk *et al.* 2010).

Fourth, changes in metrics of individual fitness like fecundity do not necessarily manifest in changes in the per-capita growth rate of the population (Ford *et al.* 2015; e.g., soil rhizobia populations often stay constant despite the >100,000-fold increase in descendants released into soil produced by a plant-associated rhizobium, Denison & Kiers 2011).

## Patterns in Historical Models

A surprising result of our work is that many of our model predictions are robust across types of mutualism and their mechanistic derivations – even among historical models (Table A1). Despite diverse approaches, many historical models predict stable coexistence and threshold effects when at least one species is obligate (May 1978, Soberón & Martinez Del Rio 1981, Wells 1983, Wolin & Lawlor 1984: Eqn. A8, Pierce & Young 1986, Wright 1989, Graves *et al.* 2006, Thompson *et al.* 2006: Eqns. A12-A13, Kang *et al.* 2011, Revilla 2015), suggesting that these outcomes could be general among mutualisms. Some historical models are even mathematically identical, despite derivations ranging from phenomenological to extremely detailed (e.g., Soberón & Martinez del Rio 1981: Eqn. A6, Wright 1989, Bazykin 1998, our Eqns. 2–3).

A theme among these historical models is that the benefits of mutualism (Fig. 4A) do not directly depend on recipient density and (Fig. 4D) are saturating, either due to interspecific density-dependence (Soberón & Martinez del Rio 1981: Eqn. A6, Dean 1983, Wright 1989, Graves *et al.* 2006, Thompson *et al.* 2006: Eqn. A12–A13 Holland & DeAngelis 2010, Kang *et al.* 2011), intraspecific density-dependence (Soberón & Martinez del Rio 1981: Eqn. A5, Wolin & Lawlor 1984: Eqn. A8, Revilla 2015: Eqn. A17), or both (May 1978, Wells 1983). Our nutrition and protection models are also in this tradition: benefits do not directly depend on recipient density and saturate due to interspecific (main text models, Eqns. 2–3) or intraspecific (alternative protection models, Appendix C) density-dependence, with qualitatively identical results (Fig. 4H).

Historical models in which mutualistic benefits directly depend on recipient density (Fig. 4B) typically describe an increase in carrying capacity due to mutualism (Whittaker 1975, Wolin & Lawlor 1984: Eqns. A6, A7, Pierce & Young 1986: Eqn. A10, Neuhauser & Fargione 2004, Thompson *et al.* 2006: Eqn. A14). Interestingly, both models with saturating benefits (Wolin & Lawlor 1984: Eqns. A6–A7) saturate without inflection (no changes in concavity) due to interspecific density-dependence (Fig. 4J). Note though that these models have been referred to as examples of *intra*-specific density dependence (e.g. Holland 2015), presumably because saturating benefits reduce negative density-dependence in the recipient population. The models predict stable coexistence (Eqns. A6–A7) or unstable coexistence (Eqn. A7, Fig. 4L), but not bistability, and can be applied to facultative mutualists only. This is in contrast to our transport models (Eqns. 4–5), which are both directly dependent on recipient density and saturate due to intraspecific density-dependence because benefits are proportional to a partner’s per-capita visitation or consumption rate on the recipient population (Fig. 4K, 4M). Additionally, high degree terms (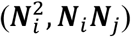, ***N***_*i*_***N***_*j*_) in their nullclines allow inflection points that can lead to bistability.

A more recent pattern in population dynamic models of mutualism investigates the stability of interactions that can shift from mutualism to parasitism, either as controlled by an interaction coefficient that increases or depresses, respectively, equilibrium density in the presence of the partner (Pierce & Young 1986, Neuhauser & Fargione 2004), or as a balance between costs and benefits that depend dynamically on partner density (Holland & DeAngelis 2010, Kang *et al.* 2011). Pierce and Young’s (1986) and Neuhauser and Fargione’s (2004) models reproduce the qualitative results of our nutrition and protection models when the interactions are mutualistic. Kang *et al.*’s (2011) model for leaf-cutter ants and their fungal gardens includes a linear cost term of fungal biomass consumption, but still reproduces the qualitative results of our nutrition and protection models between partners with Type I and Type III functional responses. Holland and DeAngelis’ (2010) models also reproduce our qualitative results when costs, which they specify are saturating costs of supplying resources to consumers, are set to zero (*q*_*i*_ = 0 in Eqn. A15). However, their full models (when *q*_*i*_ > 0) are unique because nonzero costs may exceed benefits instantaneously (“parasitism”) due to the unique dynamical outcome of overexploitation even if in the long run benefits are greater (“mutualism”). Testing for overexploitation dynamics may illuminate the prevalence of nonlinear population-level costs among mutualisms. Also see our proposed models with cost terms and comparison to Holland and DeAngelis’ work in Appendix C.

**Table A1.**
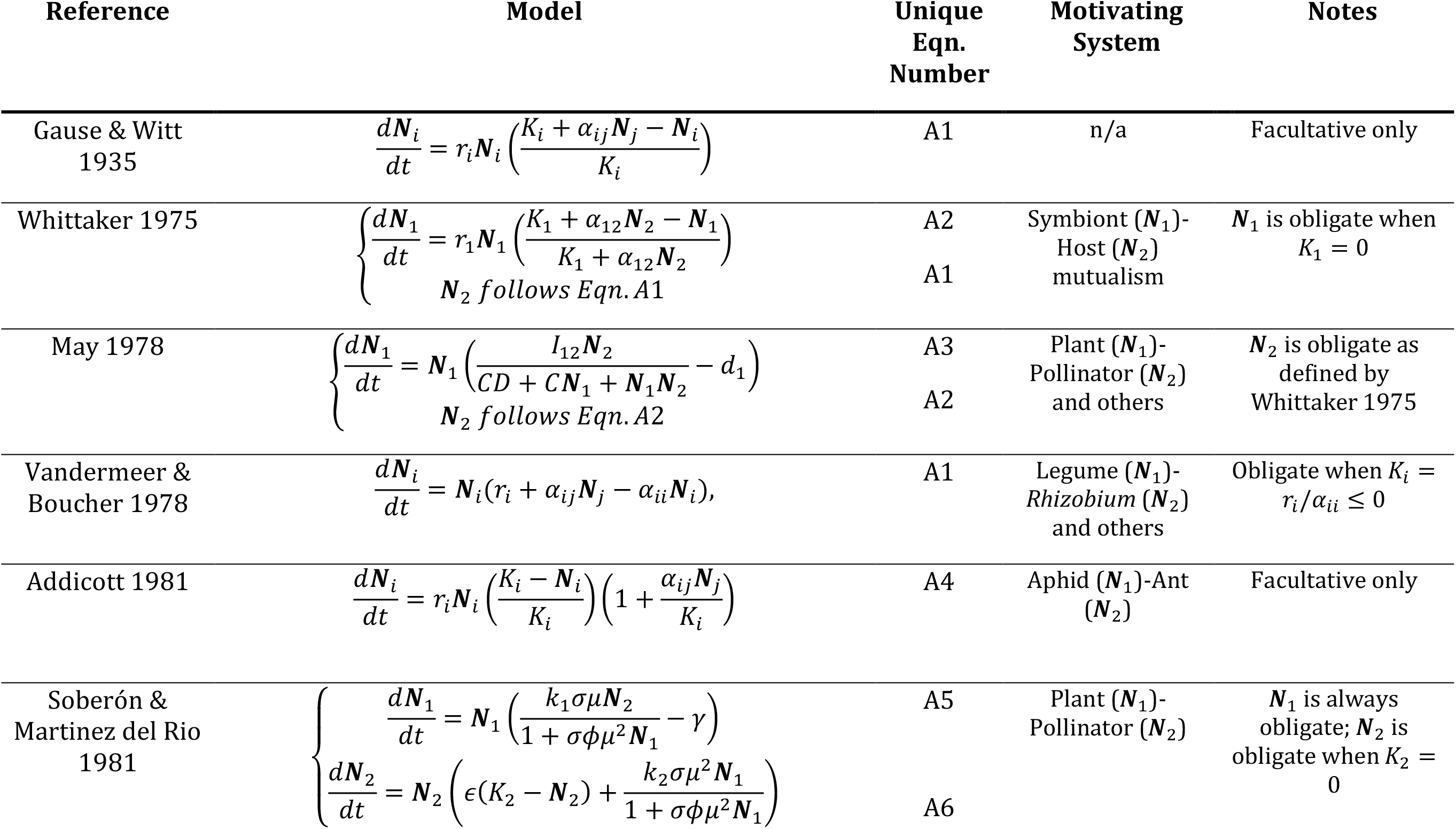
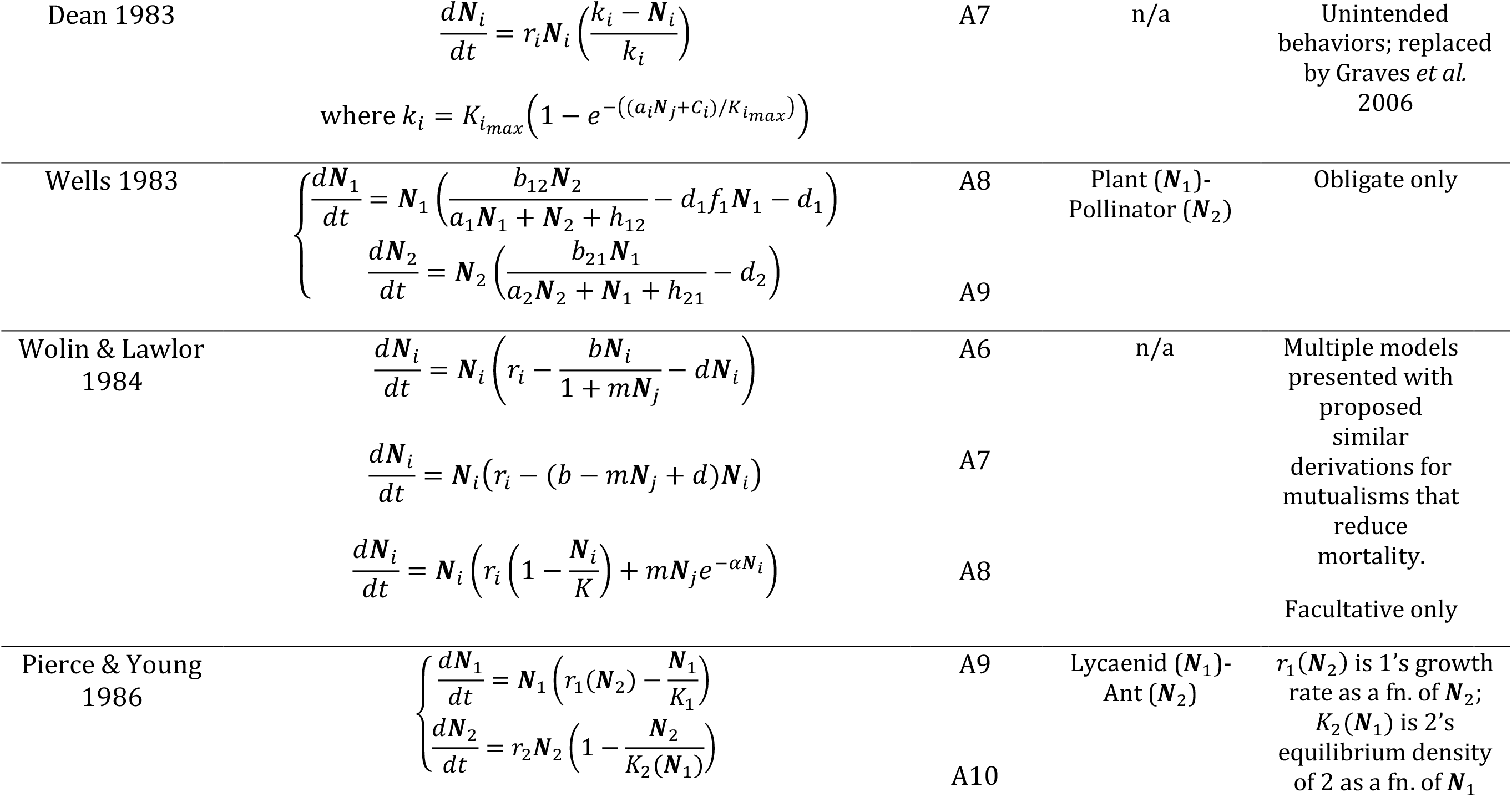
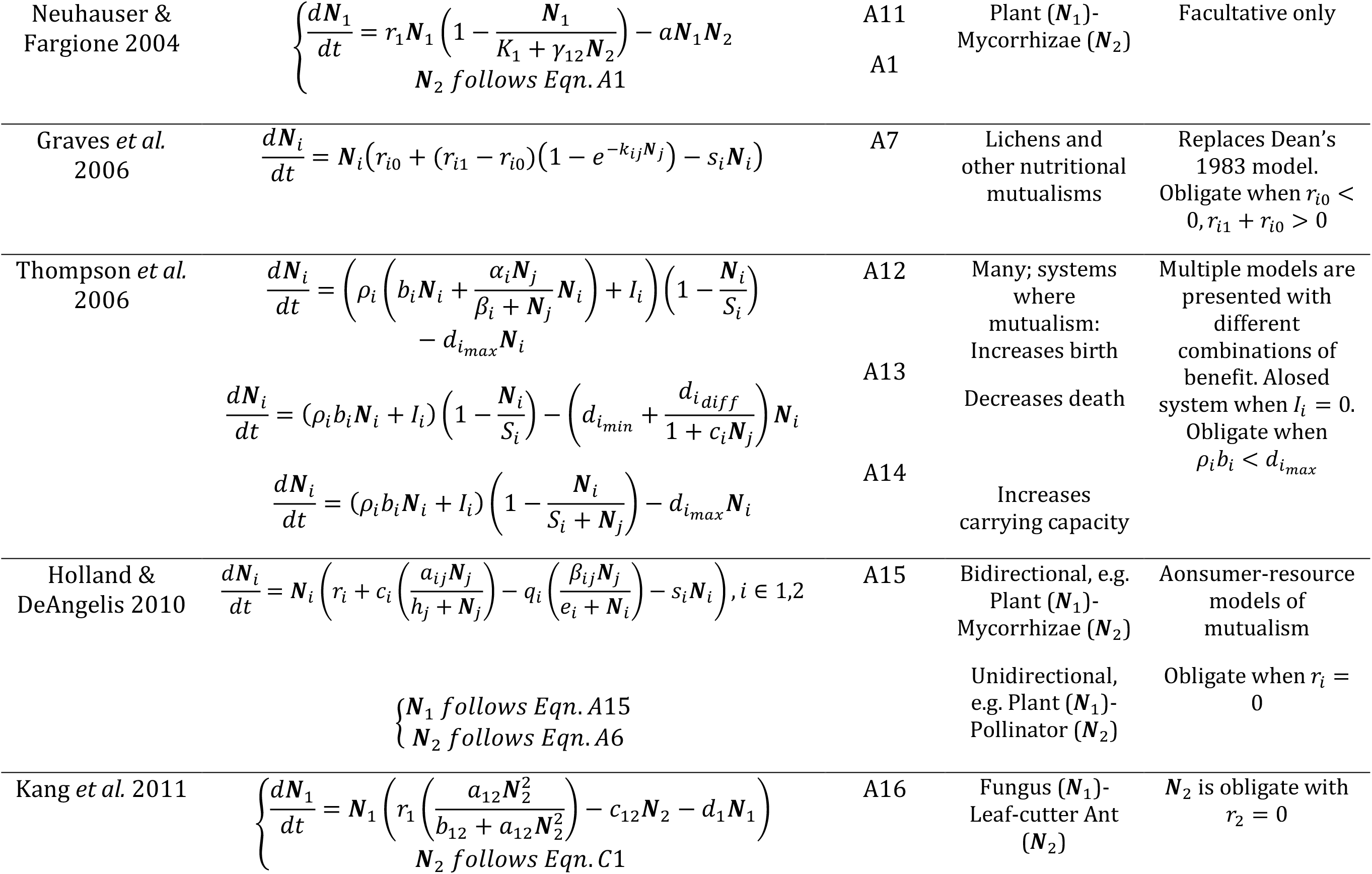
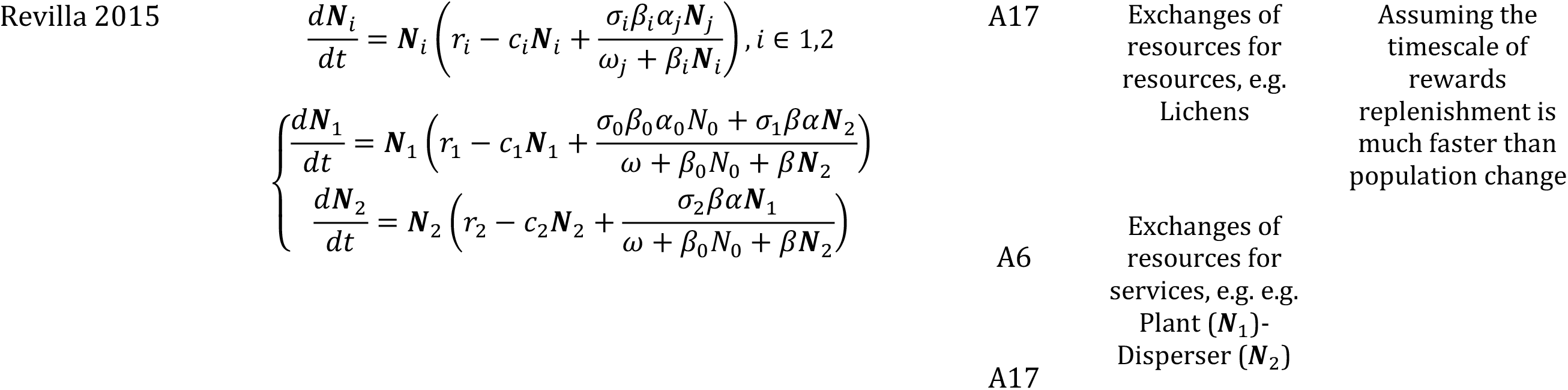
Historic population dynamic models of mutualism. This list is not comprehensive. In general, we included only a sampling of deterministic, continuous-time population dynamic models of two equations for two species that were intended by the authors to generalize across mutualistic systems. However, we also included a few specific models (e.g. Soberón & Martinez del Rio 1981, Pierce & Young 1986, Kang *et al.* 2011) if we considered their approach notable or reference their work in the text. Equations with the same numbering are not significantly different mathematically. Additionally, many models, despite different mathematical features, predict similar qualitative dynamics in the ecologically-relevant region (Appendix A). Equations largely follow the notation from the original citations. Only unique models from each citation were included. We encourage the readers to refer to the citations for the model derivations and interpretation of parameters.

## Appendix B Mathematical Details

## Analysis

We first analyze Eqn. 1 to serve as a reference for the effects of mutualism. In the absence of a potential partner (***N***_*j*_), the nullclines for Eqn. 1 are also equilibria. There are two: one “trivial”(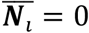, “extinction”) and one nontrivial:

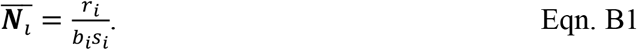

The nontrivial equilibrium is feasible (exists) when *r*_1_, = *b*_1_ − *d*_1_ ≥ 0. Because we are concerned only with dynamics when both species have non-negative density, in this work we call equilibria “stable” or “unstable” when they are attracting or repelling, respectively, in this ecological region (when ***N***_1_ ≥ 0). When *r*_1_ > 0, the trivial equilibrium is always unstable, allowing positive population growth when ***N***_1_ is at low density, and the nontrivial equilibrium is stable allowing population persistence at constant density. Specifically, when ***N***_*i*_ < *r*_1_/*b*_1_*s*_1_ population density increases due to positive intrinsic growth whereas when ***N***_*i*_ > *r*_1_/*b*_1_*s*_1_, population density decreases due to negative density-dependence.

In all of our models (Table 1), both species have “trivial” nullclines: 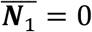, a vertical line along the y-axis, and 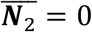, a horizontal line along the x-axis. The intersection of both species’ trivial nullclines gives a trivial extinction equilibrium (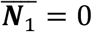,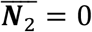) in every model. Additionally, one species ***N***_2_ follows the same dynamic equation (Eqn. 2) and thus has the same nontrivial nullclines (hereafter, simply “nullclines”). We use the Holling Type II notation in the following analysis but it is easy to substitute parameters for Michaelis-Menten kinetics. The nullcline (Figs. 1–3, gray) is a concave down, increasing function that saturates with respect to its partner’s abundance:

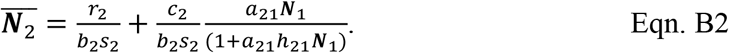

Eqn. B2 is the curve showing the balance between negative density-dependence (exerted by the (1 − *s*_*i*_,***N***_*i*_) term in Eqn. 2), benefit from mutualism, and positive intrinsic growth (when ***N***_2_ is facultative). When 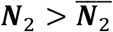, negative density-dependence is stronger than benefit from mutualism and/or intrinsic growth causing ***N***_2_ to decrease. When 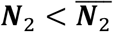, negative density-dependence is weaker and ***N***_2_ increases.

Comparing Eqns. B1 and B2, we can isolate the benefits of mutualism to ***N***_2_ as 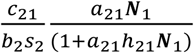. Importantly, benefits saturate with respect to partner density. In particular, as plant density becomes extremely high, ***N***_2_ saturates to a higher value than it could achieve in the absence of its partner:

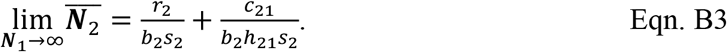

This horizontal asymptote is an upper bound on the denity ***N***_2_, When ***N***_2_ is facultative (*r*_2_ > 0) its nullcline intersects the y-axis in the ecological region (when ***N***_*i*_ ≥ 0) at

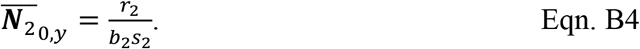

All else being equal, decreasing *r*_2_ pushes the visible part of ***N***_2_’s nullcline down so that when ***N***_2_ is obligate (*r*_2_ ≤ 0), it intersects the at y-axis at zero or negative values (not shown). Then, in the ecological region, the nullcline instead intersects the x-axis at

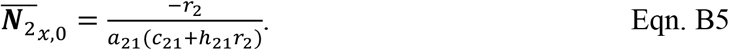

Below, we calculate the nontrivial nullclines for ***N***_1_ (Figs. 1–3, black) and similarly analyze their dynamics in the ecological region of the plane. See Methods and Tables 1–2 for terminology.

## Nutrition

Plants (***N***_1_) follow the same dynamics as microbes (***N***_2_), so their nullcline is symmetrical to Eqn. B2:

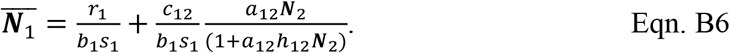

Clearly, Eqn. B6 is an increasing function that saturates with respect to microbe density. It can be rearranged to be a function of ***N***_1_:

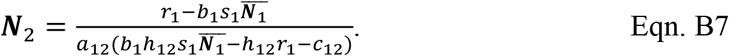

Then, plant density increases to the left of its nullcline (Fig. 1, black) due to mutualism and decreases to the right of it due to strong negative density-dependence. All other properties are likewise symmetrical (i.e. Eqns. B3–B5 apply with the M and N axes switched and indices *x* and *y* swapped).

## Protection

Species that benefit from protection mutualisms (***N***_1_) follow the same dynamics as nutritional mutualists. In the protection notation, their nontrivial nullcline (Fig. 1, black) is:

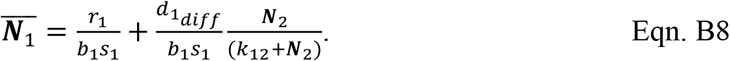

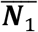 approaches a vertical asymptote at

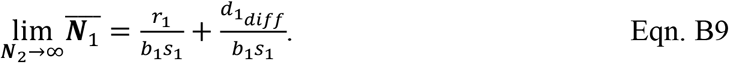

When obligate (*r*_1_ < 0), 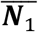 intersects the y-axis in at

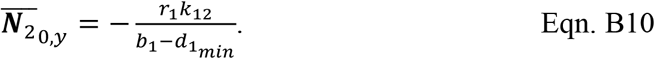

When facultative (*r*_*1*_ > 0), 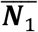 intersects the x-axis at

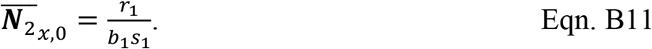

## Transport

The equation for the nullcline of animal-pollinated plants (Fig. 2, black) is:

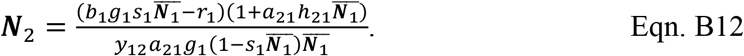

When plants are obligate (*r*_*1*_ = *g*_*1*_*b*_*1*_ − *d*_*1*_ ≤ 0, Fig. 2a-b), the nullcline is a top-open hump bounded by vertical asymptotes at ***N***_1_ = 0 and ***N***_1_ = 1/*s*_1_. When plants are facultative (*r*_*1*_ > 0, Fig. 2c-g), at high density the nullcline is increasing, concave up, and saturates at 1/*s*_1_. At low density, the nullcline is concave down, though this may not be visible in the ecological region of the plane. Regardless, it has x-intercept

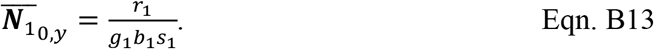

The equation for the nullcline of animal-dispersed plants (Fig. 3, black) is:

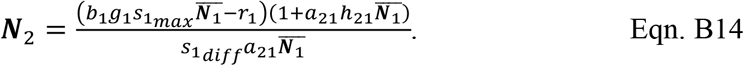

At high plant density, the nullcline has a linear slope of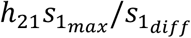, which is ≥ 1 due to our condition that negative density-dependence can be reduced at most to zero (see Methods). When plants are obligate (*r*_*1*_ = *g*_*1*_*b*_*1*_−*d*_*1*_ ≤ 0, Fig. 3a-b), the nullcline is a top-open hump bounded on the left side by a vertical asymptote at ***N***_1_ = 0. When plants are facultative (*r*_1_ > 0, Fig. 3c-g), the nullcline is concave down at low density, with x-intercept

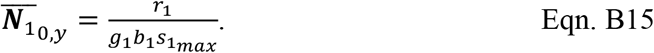

For both Eqns. B12 and B14, inside the hump or to the left of the curve plant density increases; to the right, plant density decreases due to strong negative density-dependence.

## Feasibility

All of our models are structurally unstable (Rohr et al. 2014), such that smooth transitions in parameter values shift the nontrivial nullclines (hereafter, “nullclines”) so that they may intersect in various ways or even fail to intersect, with different dynamical outcomes for each case (Fig. B1). In the main text, we focused on “feasible” systems, that is, where coexistence of the partners with non-negative density (i.e. in the ecological region) was possible. There are many parameterizations that lead to feasibility, but it is not easy to provide an ecological interpretation for the conditions to achieve them. Below we enumerate the geometric conditions that lead to feasibility by describing the required positioning of the nullclines.

## Nutrition and Protection

There are three cases for how the nullclines of ***N***_1_ and ***N***_2_ intersect:

1. The nullclines never intersect (Fig. B1a). Coexistence is not possible. Depending on the initial conditions, both species may be extinct or a facultative species may persist at 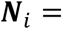 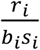 with ***N***_*j*_ = 0.
2. The nullclines intersect exactly once (Fig. B1b). This case is extremely sensitive to the vital rates like birth and death rates of both species; it is highly unlikely in nature these rates would be maintained at such precise values, especially in the presence of stochasticity. We neglect this case.
3. The nullclines intersect twice (Fig. B1c-d), yielding exactly one saddle point at low density and one stable equilibrium at high density. Coexistence is feasible when at least one of the intersections is in the ecological region (Fig. B1c, d: two, one intersection(s) in the ecological region, respectively). When only one intersection occurs in the ecological region, it is the stable coexistence equilibrium.

## Transport

There are four cases for how the plant and animal nullclines intersect:

1. If the nullclines never intersect (Fig. B1e), then coexistence is not possible.
2. If the nullclines intersect exactly once when plants are obligate (Fig. B1f), this is an ecologically unlikely case that we neglect.
3. If the nullclines intersect twice with at least one intersection in the ecological region (e.g. Fig. B1g, B1h), then coexistence is feasible. The low-density equilibrium is a saddle point while the high-density equilibrium is stable. When only one intersection occurs in the ecological region, it is the stable coexistence equilibrium.
4. If the nullclines intersect three times in the ecological region (e.g. Fig. B1i), then coexistence is feasible. The low and high density equilibria are stable (“bistability”) and the intermediate equilibrium is a saddle point. Bistability appears to only occur when at least one nullcline changes concavity in the ecological region. Our transport models display bistability because their nullclines change from concave up to concave down (pollination, Figs. 2d, 2f) or from concave down to linear growth (seed dispersal, Figs. 3d, 3f). However, bistability can more generally result from Type III functional responses in facultative mutualists (see Appendix C).

**Figure B1.**
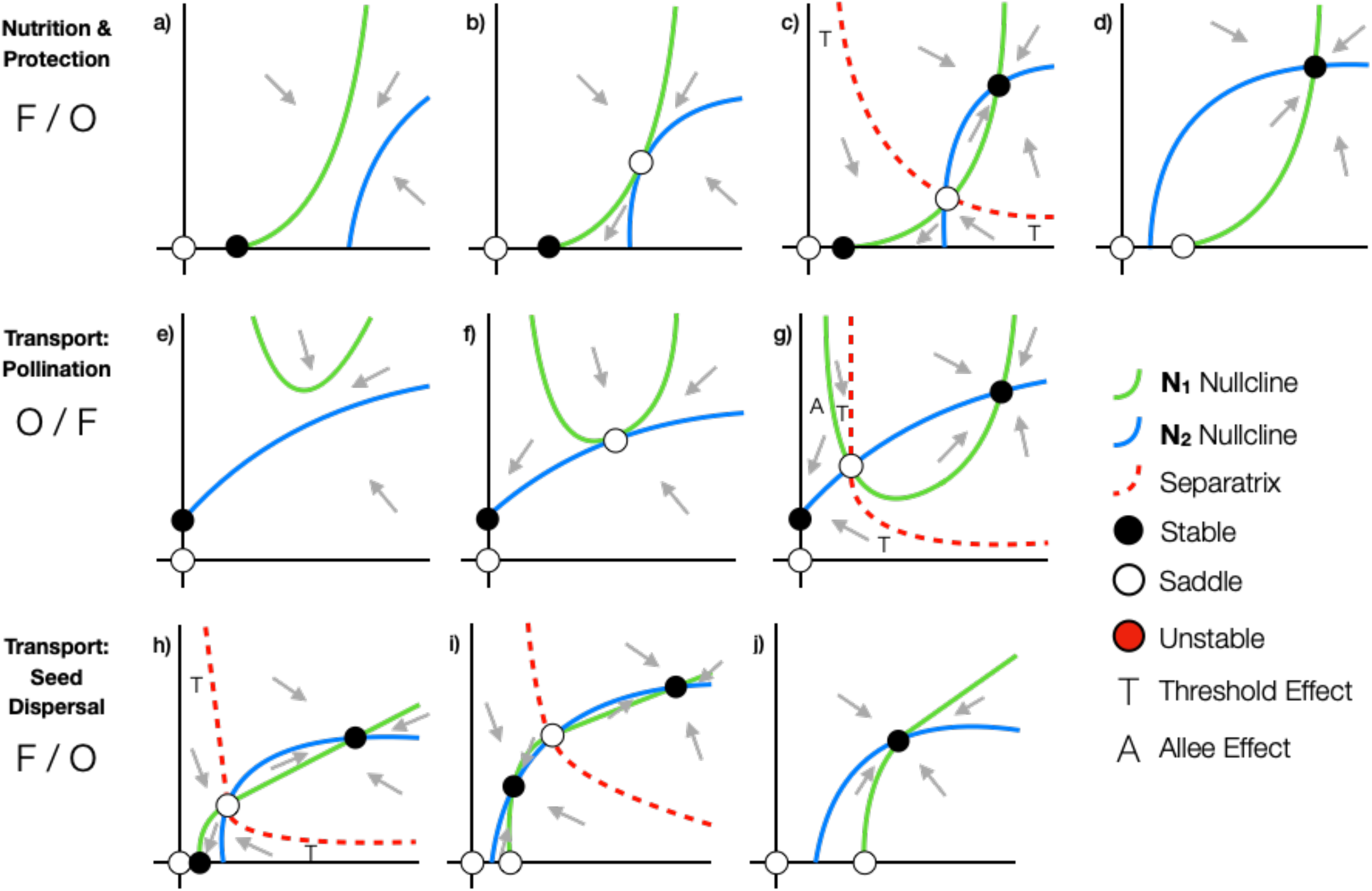
Examples of structural instability. Phase plane diagrams illustrating smooth transitions from infeasible to feasible coexistence (from left to right panels within a row). Formatting follows Figs. 1–3. Labels for each row identify the models being illustrated (Table 1) and species 1 / species 2 as obligate (O) or facultative (F). (**a-d**) The ***N***_2_ nullcline shifts smoothly to the left, due to, for example, increasing conversion efficiency (***C***_21_). This moves the system from (**a**) infeasible, to (**b**) ecologically unlikely, through a “blue sky” bifurcation in which (**c**) coexistence is feasible and stable but will not occur if species’ initial densities fall below a threshold, to (**d**) guaranteed stable coexistence. This type of structural instability occurs in both our nutrition and protection models (shown) and in our pollination model when plants are facultative (not shown). (**e-g**) the vertex of the ***N***_1_ nullcline shifts smoothly down, due to, for example, increasing partner quality (*y*_12_). This moves the system from (**e**) infeasible, to (**f**) ecologically unlikely, through a blue sky bifurcation in which (**g**) coexistence is feasible and stable but threshold and Allee effects are present. (**h-j**) The ***N***_1_ nullcline shifts smoothly to the right, due to, for example, decreasing maximum negative density-dependence 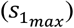, i.e. negative density-dependence in the absence of the partner. This moves the system from infeasible (not shown), to ecologically unlikely (not shown), through a blue sky bifurcation in which (**h**) coexistence is feasible and stable but threshold effects are present, to (**i**) bistability in which stable coexistence is guaranteed but occurs either at a low- or high-density equilibrium, to (**j**) guaranteed stable coexistence. This type of structural instability occurs in both our seed dispersal model (shown) and in our pollination model when plants are facultative (not shown).

## Appendix C

## Functional Forms

To understand the robustness of our results, we analyzed our models (Table 1) using other functional forms in addition to the saturating forms presented in the main text. Specifically, we report all possible dynamical outcomes for every combination of obligate and facultative partners with Holling Type I, II, and III functional responses. Table C1 specifies the mathematical expressions used for each model. Notice that in the transport models, reproductive services (*S*) to population *i* are a function of partner *j*’s consumption or visitation rate (*C*_*R*_) on *i*, so only the functional form for *j* needs to be specified.

Our results are generally robust to variation in functional form (Figs. C1–C3). We highlight the exceptions to that pattern here. In our nutrition and protection models (Fig. C1), coexistence is infeasible or leads to the orgy of mutualism (that is, unstable coexistence) when both partners have Type I functional responses. Additionally, bistability becomes a possible dynamical outcome when a facultative partner has a Type III functional response. In our pollination model (Fig. C2), all dynamical outcomes are as reported in the main text, even when pollinators follow a Type I functional response. In our seed dispersal model (Fig. C3), the orgy of mutualism always occurs when dispersers follow a Type I functional response.

Though we do not analyze them here, other functional forms including the Beddington-DeAngelis formula, ratio-dependent forms, and unimodal forms should be used in our models according to the specific system (reviewed by Holland 2015). Note that for nutrition and transport mutualisms, we derived models in which the functional responses (or more precisely, numerical responses, Revilla 2015) are direct functions of recipient or partner consumption rate. However, our models can also accommodate functional responses interpreted as “net benefit” curves (e.g., Holland *et al.* 2002, Morris *et al.* 2010) if costs and benefits affect the same vital rate (i.e., if “costs” simply reduce the benefits that accrue to a given vital rate). Under this interpretation, unimodal functional forms may arise (e.g. Morris *et al.* 2010) which could lead to substantially different dynamical predictions than those presented here.

## Alternative Models

In the main text, we presented a protection model for species *i* in which the partner population (***N***_*i*_) reduces mortality (*d*)) by deterring natural enemies (Eqn. 3). If *j* instead reduces mortality via attendance on the recipient population (e.g. Morales 2000), we choose 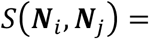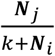 or 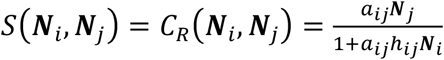 so that per-capita death rate declines proportional to *j*’s recruitment rate or consumption rate on *j*’s rewards, respectively. Then, benefit saturates when *j*’s density is high. Here, we analyze the specific model:

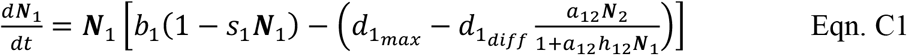

where attendees (***N***_1_) are obligate on protectors (***N***_2_) when 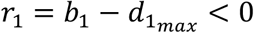 and facultative otherwise. We additionally require 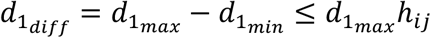, so that net mortality is always non-negative.

Alternatively, we could choose the simplest function so that mortality is reduced monotonically with saturation when partner density is high, as proposed by Thompson *et al.* (2006). In this case, we modify Eqn. 3 to give the specific model:

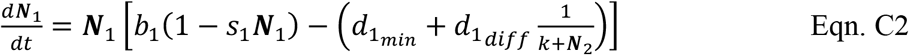

where, again, attendees (***N***_1_) are obligate on protectors (***N***_2_) when 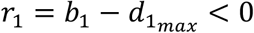 and facultative otherwise.

Another alternative is to assume that protection services increase attendees’ maturation rate (*b*) as in the case where ants (***N***_2_) guard the pupae of lycaenid butterflies (***N***_2_, Travassos & Pierce 2000). Then, we modify Eqn. 3 to give the specific model:

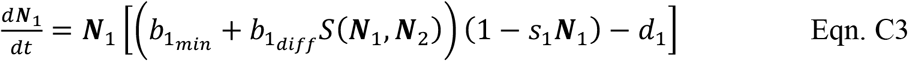

where we choose 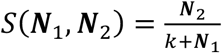 to represent ant recruitment rate. Lycaenids are obligate on ants when 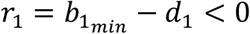 and facultative otherwise.

Finally, we could assume that tending by protectors decrease attendees’ mortality due to density-dependent processes (*s*), which may act over the course of attendees’ lives. For example, ants reduce aphid mortality through cleaning and quarantining practices that (presumably) prevent the spread of fungal infections (Nielsen *et al.* 2010, Durak *et al.* 2016). Then, we modify Eqn. 3 to give the specific model:

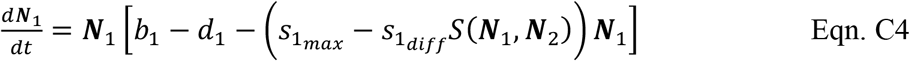

where we again choose 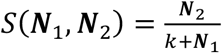 to represent ant recruitment rate. Aphids (***N***_1_) are obligate on ants (***N***_2_) when *r*_1_ = *b*_*i*_ − *d*_*i*_ < 0 and facultative otherwise.

We can also consider an alternative model for seed dispersal. In the main text, we presented a model (Eqn. 5) wherein plants benefit from seed dispersal by escaping density-dependent sources of mortality during maturation (*S*). Some plants (***N***_1_) instead benefit exclusively via an increase in seed survival/germination rate (*s*) because gut passage through avian or mammal dispersers (***N***_2_) removes pathogens from seeds and provides chemical camouflage from seed predators (Fricke *et al.* 2013). In this case, we modify Eqn. 5 to give the specific model:

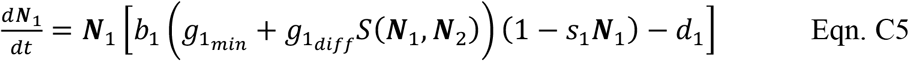

where plants are obligate on dispersers when 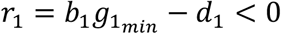 and facultative otherwise. We choose 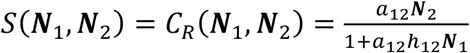 to represent the consumption rate of fruit by dispersers. Since we originally defined *g* as the fraction of seeds that germinate, we additionally require 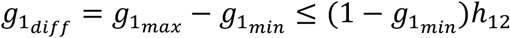, so that net germination fraction is always between 0 and 1.

Protection models C1-C3 have similar qualitatively nullcline geometries as our main text protection model (Fig. C4a-d). Though the nullcline for model C1 does not saturate (Fig. C4b), it will yield the same dynamical outcomes as our main text model (Fig. 2) when paired with a partner such as a nutritional mutualist with a concave down saturating or linear nullcline. Models C2-C3 have exactly the same qualitative nullcline geometries as our main text model, despite mutualism potentially affecting different vital rates in these models (Fig. C4c-d). In all cases (Eqns. C1–C3), as in our main text model (Eqn. 3), the effect of mutualism is independent of recipient density.

In contrast, protection model C4 has qualitatively identical nullcline geometry to our main text seed dispersal model (Fig. C4e), because in both cases the effect of mutualism is dependent on and saturating in terms of recipient density. In model C4, protection services affect a density-dependent process (modifying *s*) and benefit is proportional to attendance or recruitment rate by protectors on the recipient species (saturates via intraspecific density-dependence).

Finally, seed dispersal model C5 has qualitatively similar nullcline geometry to our main text nutrition and protection models (and to protection models C1-C3), because the benefits of seed dispersal (increased germination) are not directly dependent on recipient density.

## Extensions

The extensions proposed here are not mutually exclusive and can be applied to any of our main text models. First, we consider an extension that accounts for the costs of rewards exploitation, including the special case in which depletion of rewards due to exploitation induces variable and potentially high costs of rewards production. Exploitation costs are those incurred while a partner is acquiring rewards, such as damage to flowers by pollinators during foraging for nectar and pollen. Such costs accumulate proportionally to the exploiter’s visitation or rewards consumption rate, which can affect individual fitness and demographic rates, and is often assumed to affect population growth rate as well (e.g. Aizen *et al.* 2014). Interestingly, rewards production costs can also accumulate proportionally to rewards consumption rate. For example, when tended by ants, some aphid species enrich their honeydew with synthesized sugars, which decreases their fecundity and body size, presumably due to carbohydrate depletion (Yao *et al.* 2000, Yao & Akimoto 2001). Though honeydew enrichment is a construction cost (rewards production), it can be modeled as proportional to rewards consumption (exploitation) because ants’ consumption behavior induces the metabolic costs of enriching depleted honeydew.

For easy comparison to previous works that include costs (e.g. Holland & DeAngelis 2010, Table A1), we modify our nutritional model with Michaelis-Menten functional responses to include rewards exploitation costs as follows:

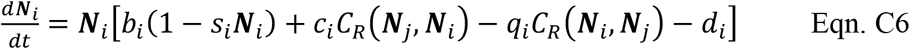

where costs to population *i* are related to the benefits to population # because they occur via the same mechanism: *j*’s exploitation of *j*’s rewards. Mathematically, this is a special case of Holland and DeAngelis’ (2010) model (specifically, our model has four fewer parameters), which assumed independently parameterized functional responses for benefits and costs. The coefficient *q*_1_ controls the effect of rewards exploitation on population density. If rewards exploitation has minimal population-level effects, 0 ≤ *q*_1_ ≪ 1. Assuming both species incur exploitation costs yields the specific model:

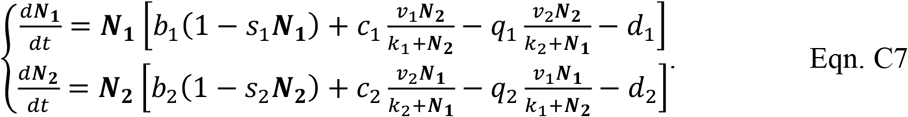

Our simple nutrition model (Eqn. 2) yields an increasing, concave down nullcline that saturates with respect to the partner’s density. Including an explicit term for saturating costs of rewards consumption (Eqn. C7) bends species’ nullcline towards the partner’s axis at high partner density, eventually curving it back around towards the origin into a lobe shape (Fig. C5a). This is because high partner density exerts high saturating costs on the recipient due to rewards consumption, which may exceed the benefits that can be acquired. When coexistence is feasible, up to three non-trivial equilibria occur: a stable node flanked by two saddle points. Stable coexistence can occur at higher densities than either species could achieve alone. However, separatrices running through the saddle points define basins of attraction that lead to extinction or, if at least one species is facultative, single-species persistence. This ensures instability when one population is of substantially higher density than the other due to overexploitation of the rare partner (regions labeled “E,” Fig. C5a). This is contrast to threshold effects defined in the main text wherein the low-density partner benefits from mutualism but cannot provide sufficient reciprocal services. When the low-density partner becomes even rarer, it experiences an Allee effect, leading to its extinction (“A,” Fig. C35a). The high-density partner will also go extinct if it is obligate upon the low-density partner. See Holland and DeAngelis (2010) for a complete analysis and Cropp and Norbury (2018) for a summary of the model’s behavior.

Second, we consider an extension that accounts for dynamics of consumption of individuals in a partner population. For example, butterflies and moths often act as herbivores as larvae (damaging or killing plant vegetation, *V*) and as pollinators (consuming nectar rewards, *R*) when mature. Similarly, protector ants sometimes consume individuals of their attendee population (aphids, lycaenids, plants, etc.) in addition to the provided rewards (honeydew, nectar, etc.). This ‘vegetative’ consumption (*C*_*V*_(***N***_*i*_)) directly reduces the attendees’ density:

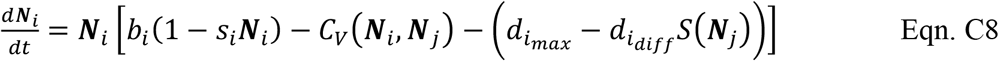

and increases the protectors’ density:

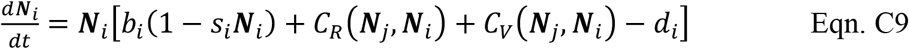

Specifically, we assume that rewards density is proportional to the attendees’ density (***N***_1_), and that ants (***N***_2_) forage according to a Holling Type II functional response on both rewards and ***N***_1_ (attendee individuals) but with different attack rates (*a*_2*R*_, *a*_2*y*_) and handling times (*h*_2*R*_, *h*_2*y*_):

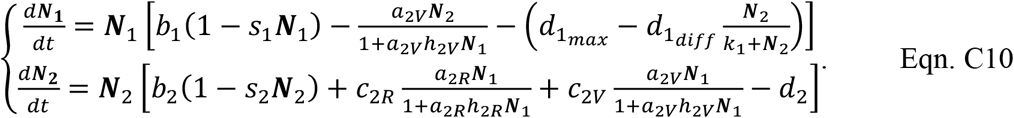

Then, attendees exhibit dynamics of an identical mathematical form to nutritional mutualists with rewards exploitation costs (Eqn. C6 with *q* = 1 indicating consumption of individuals in the population). Protectors that access both rewards and individuals from their partner population exhibit qualitatively similar dynamics to protectors that consume only rewards (Eqn. 2): the nullclines are concave down, increasing curves that saturate with respect to attendee density. However, accessing an additional resource allows the protector population to saturate to a higher density (of 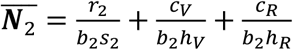) than could be supported by rewards alone.

In our extended model (Eqn. C10), protector and attendee nullclines may intersect once or twice with similar dynamical outcomes as in our simple model (Fig. 1). When the nullclines intersect exactly twice, the lower-density equilibrium is a saddle point that induces a separatrix, under which overexploitation of attendees by protectors destabilizes the system by inducing an Allee effect in the attendee population that causes extinction of facultative partners. Our simpler model also exhibits destabilization that can result in extinction, but due to threshold effects instead of overexploitation (Fig. 1A, B, D).

Our extended model also allows for the protector and attendee to intersect three times, leading to unique dynamical outcomes (Fig. C5b). Here, a separatrix divides the phase plane into two regions. On the right side, a basin of attraction allows stable coexistence to be maintained at a single high-density node (non-oscillatory coexistence). On the left side, overexploitation (“E,” Fig. C5b) by the protector population causes an Allee effect (“A”) in the attendee population, which does not necessarily lead to extinction. After depleting their resource population, the protector population also declines, eventually allowing the attendee population to receive sufficient benefit via protection compared to losses due to consumption. The system thus recovers and coexistence is maintained in this region via a limit cycle (i.e. oscillations) around a stable center, an outcome not seen in our simpler models.

**Table C1.**
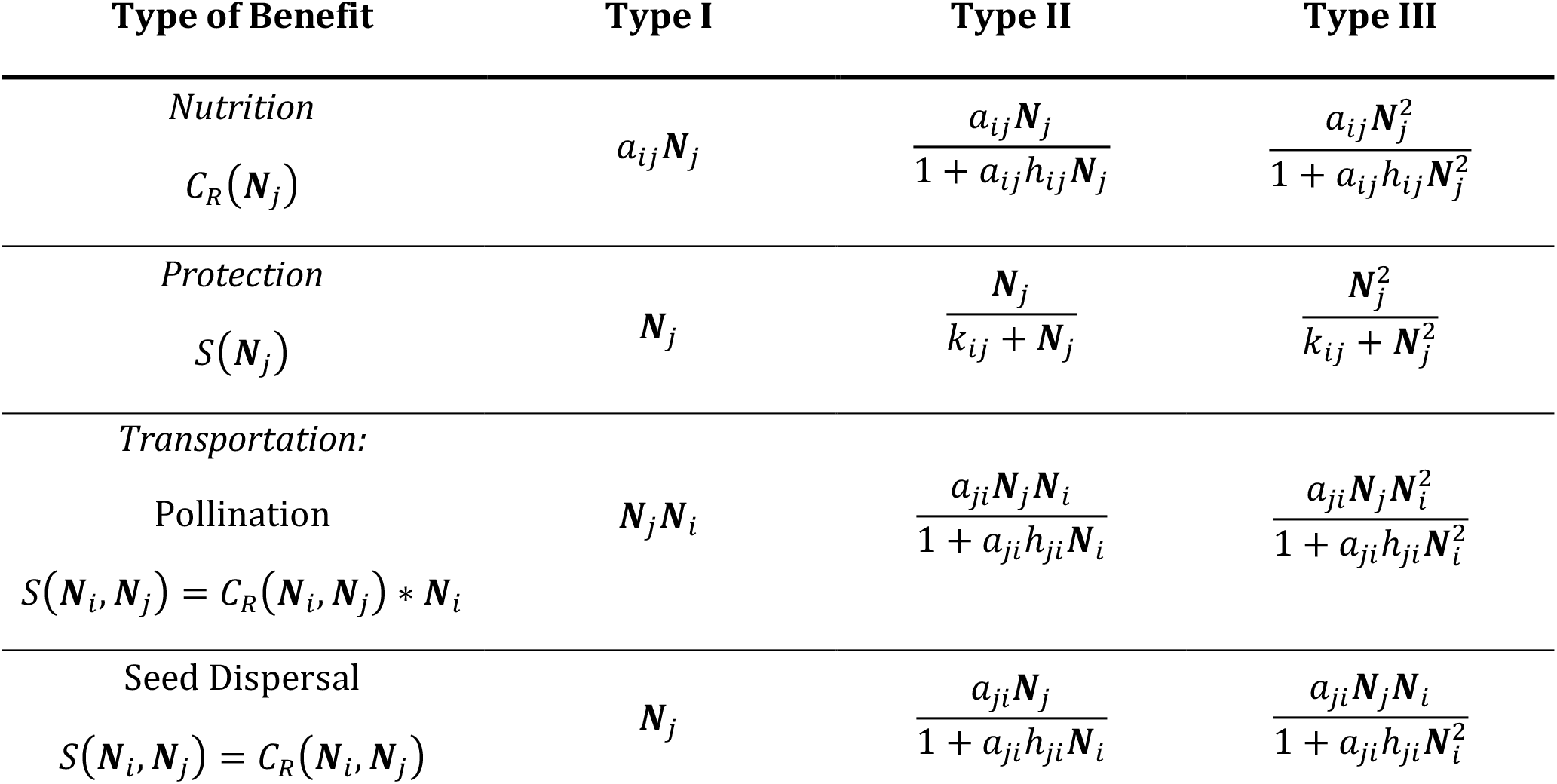
Functional forms. Functional forms used to assess robustness of results for each model. Functional forms describe the consumption rate on a partner’s rewards (*C*_*R*_) or the accrual of services provided by a partner (*S*), which, in the transportation models, are a function of the partner’s consumption rate. Indices *i* and *j* refer to the recipient and partner population, respectively. For the nutrition model, we use the Holling notation for convenience.

**Figure C1.**
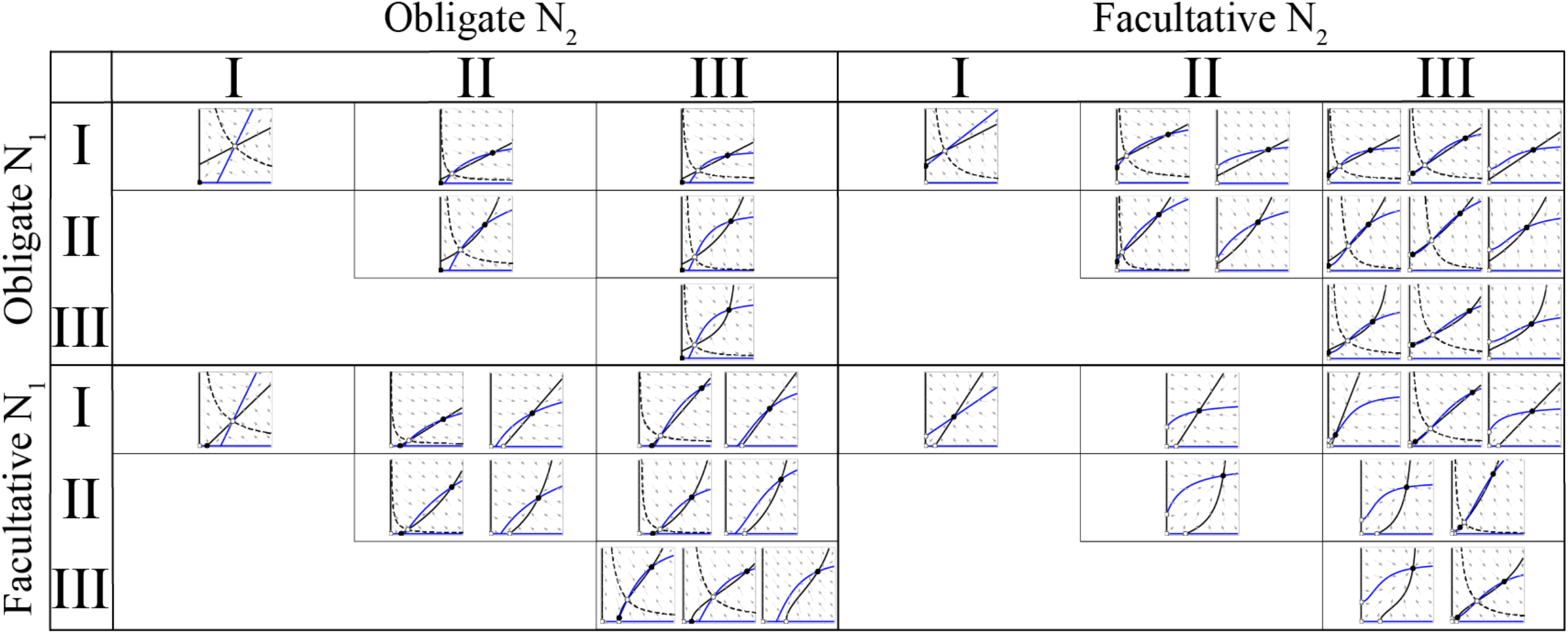
Effect of functional form on nutrition and protection mutualisms. Panels show phase plane diagrams with unique dynamical outcomes when each partner population (***N***_1_, black; ***N***_2_, blue) is an obligate or facultative mutualist following a Holling Type I, II, or III functional response (Table C1). Equilibria are stable (filled, black) or unstable or saddle points (hollow). Saddle points are bisected by a separatrix (dashed black line) that divides the plane into two regions.

**Figure C2.**
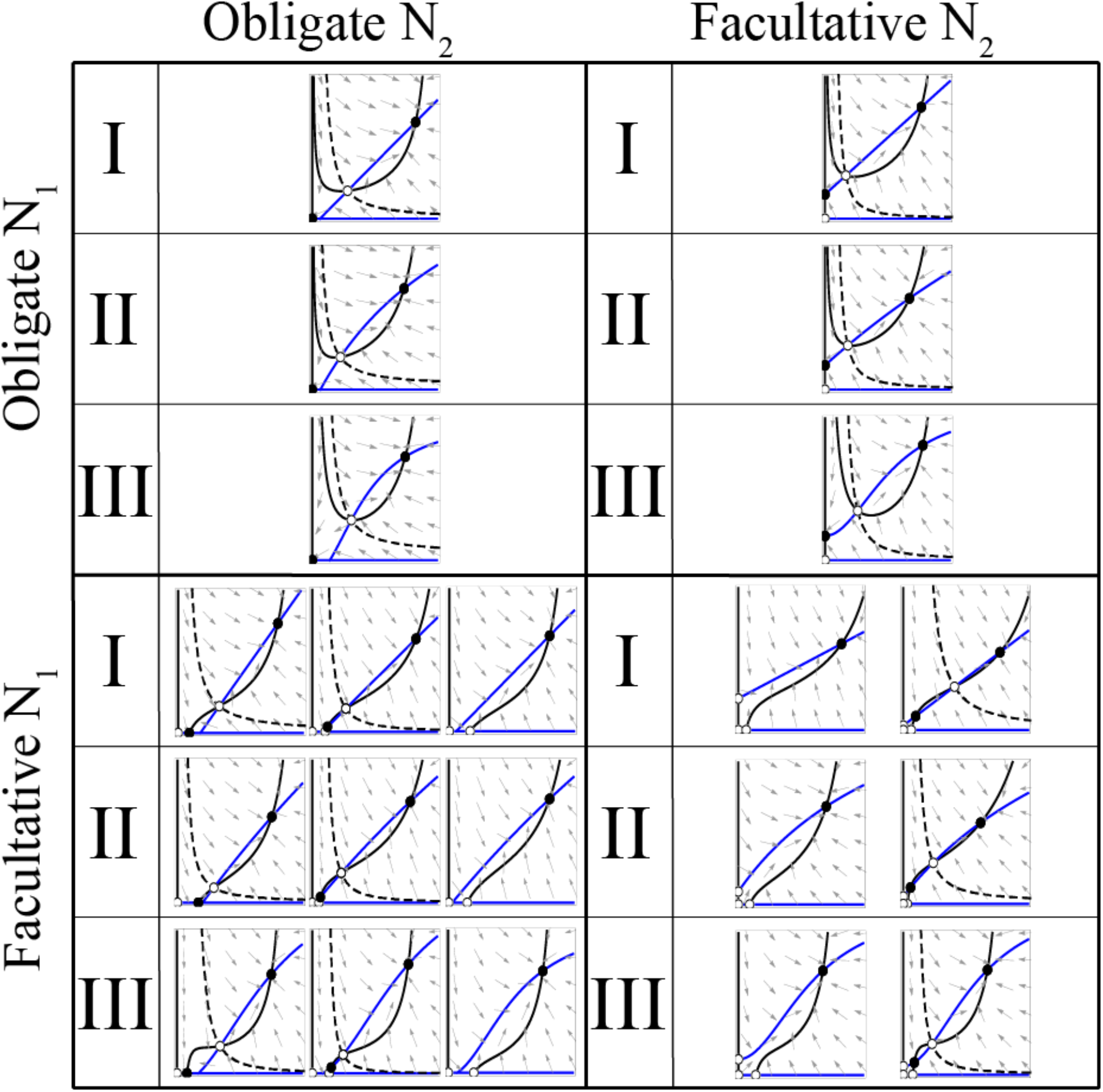
Effect of functional form on pollination mutualisms. Formatting follows Figure C1. Plants (***N***_1_) accumulate benefit proportionally to the functional form of animal pollinators (***N***_2_).

**Figure C3.**
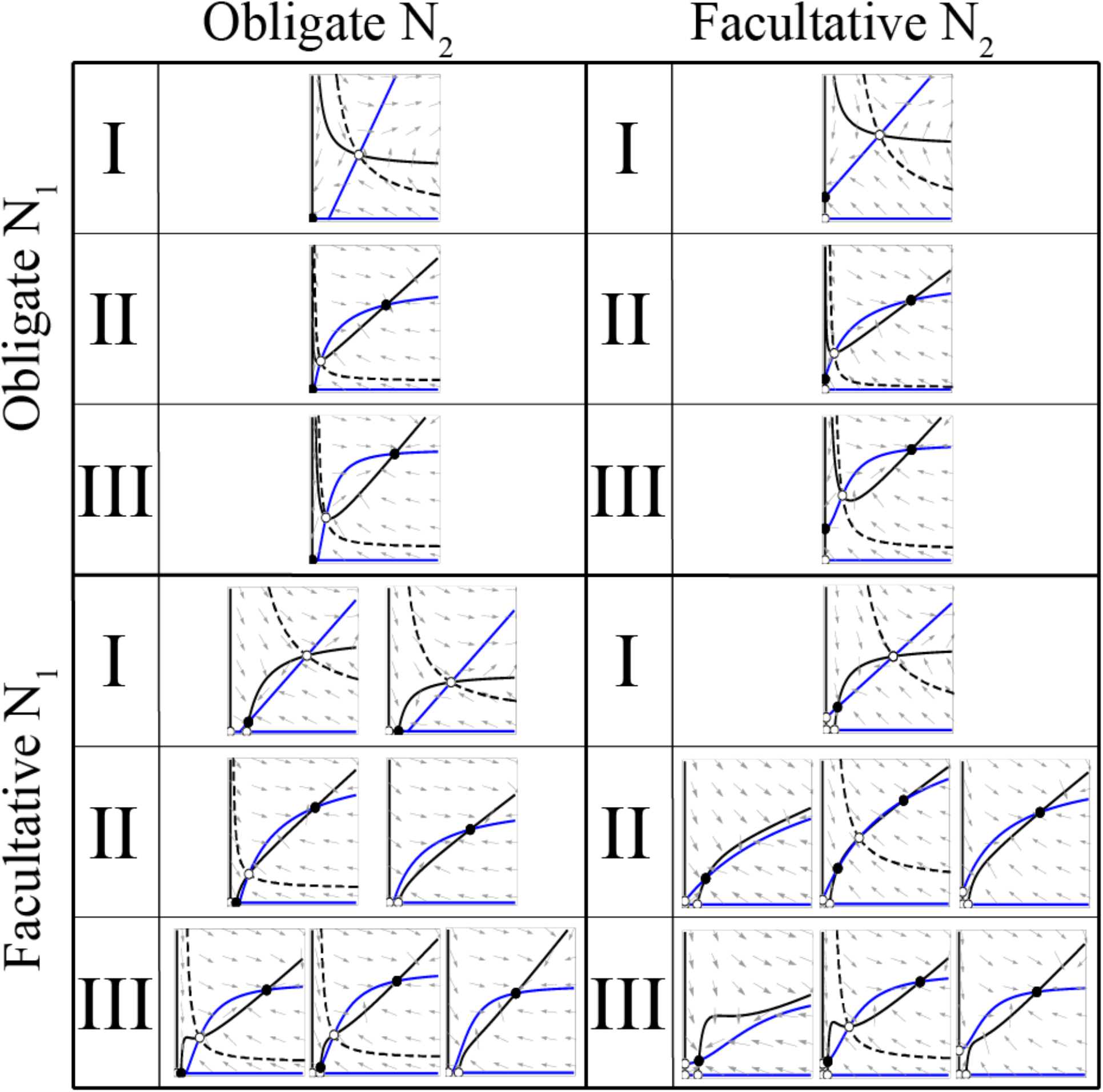
Effect of functional form on seed dispersal mutualisms. Formatting follows Figure C1. Plants (***N***_1_) accumulate benefit proportionally to the functional form of animal pollinators (***N***_2_).

**Figure C4.**
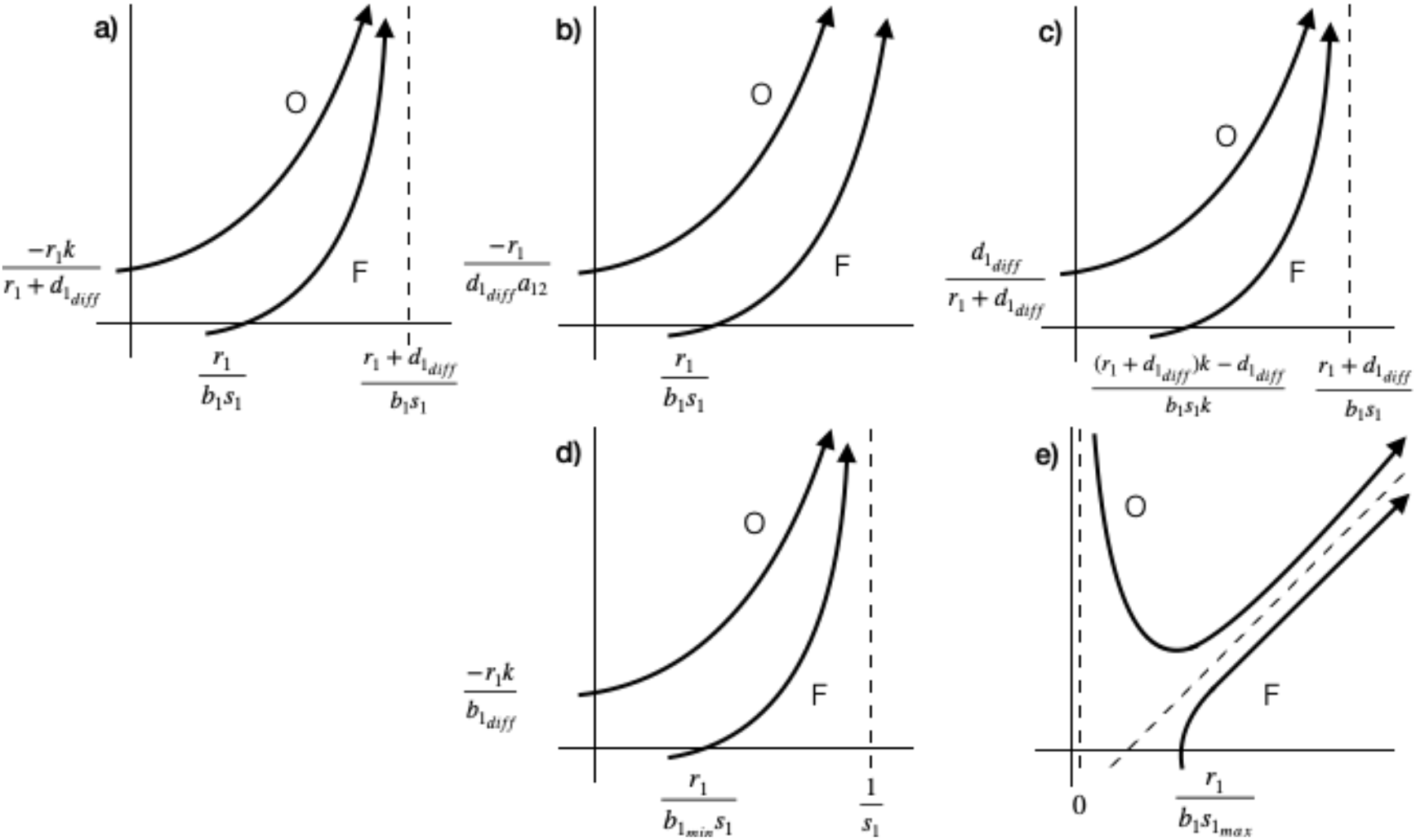
Nullcline geometry of alternative protection models. Only geometries when coexistence is feasible are shown. Nullclines (black lines) intersect the x-axis when recipients of protection services are facultative (F) mutualists or the y-axis when they are obligate (O). Asymptotes are shown with dashed lines. (**a**) Main text model (Eqn. 3). (**b**) Model C1. (**c**) Model C2. (**d**) Model C3. (**e**) Model C4. (**b**-**d**) Benefits are not directly dependent on recipient density, but saturate due to (**c**) inter- or (**b**, **d**) intraspecific density-dependence. (**d**) Benefits are additionally limited by negative density-dependence. (**e**) Benefits are directly dependent on recipient density and saturate due to intraspecific density-dependence.

**Figure C5.**
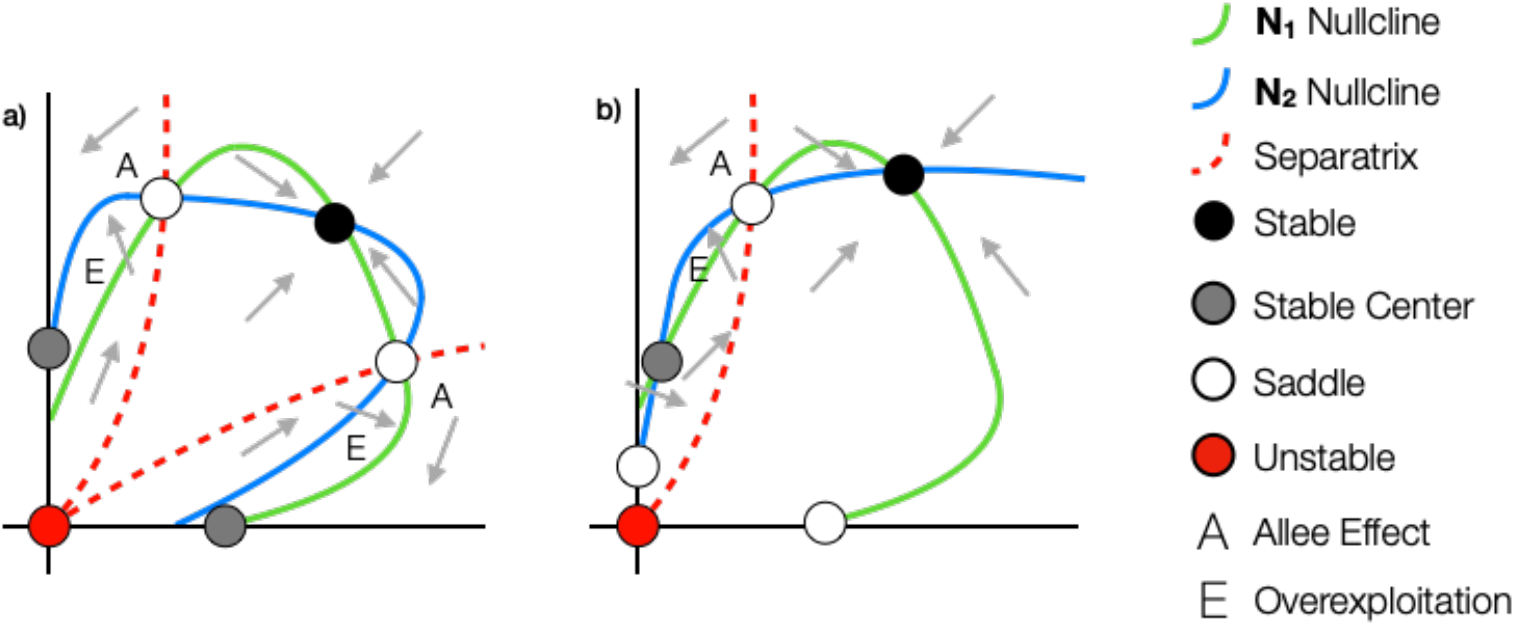
Phase plane diagrams for our extensions. Only a sampling of cases is shown for comparison to our main text models (Figs. 1–3) and Holland and DeAngelis’ (2010) models including cost terms (Table A1). (**a**) Model C7. (**b**) Model C10.

